# Defining reversible binding rates in 1D systems dependent on diffusion, density, and excluded volume

**DOI:** 10.64898/2026.06.18.733157

**Authors:** Mankun Sang, Margaret E. Johnson

## Abstract

Binding reactions in effectively one-dimensional systems, such as proteins diffusing along DNA or other filaments, pose a fundamental coarse-graining challenge because stochastic trajectories are recurrent in one dimension and therefore do not admit a unique, separation-independent macroscopic association rate. As a result, continuum rate equations are not exact in 1D even for initially homogeneous systems. Here we develop a practical framework for mapping stochastic 1D reaction-diffusion dynamics onto effective kinetic models. Using mean-first-passage arguments and particle-based simulations, we define a density-dependent association rate and a corresponding single-rate approximation, and quantify when each provides an accurate description of the underlying stochastic dynamics. We implement 1D reaction-diffusion with excluded volume in the NERDSS software using a free-propagator reweighting algorithm and validate it against known pairwise and many-body limits. Our results show that ordinary rate equations with a single effective rate can accurately reproduce 1D reaction kinetics when the dimensionless parameter governing the ratio of intrinsic to diffusion-limited reactivity is small, with excellent agreement in the strongly rate-limited regime and increasing deviations as diffusion control strengthens. We further show that excluded volume in 1D can appreciably alter both kinetics and equilibrium populations, even at modest particle densities, by reducing accessible length and introducing blockade effects. Together, these results provide quantitative guidance for selecting between spatial simulations, density-dependent rate models, and single-rate continuum descriptions of reversible 1D binding reactions.

## Introduction

Stochastic binding interactions between species restricted to 1D diffusion are recognized as fundamental steps in essential biological processes like DNA transcription ^1,2^, and transport along cytoskeletal filaments ^3–5^. A well-known application is the binding of a protein to a specific DNA sequence motif via 1D sliding along the DNA ^6^. Combining this 1D association with 3D association, or facilitated diffusion^7^, has been extensively characterized via both theory ^6,8–12^ and experiment ^13–17^. However, proteins can perform 1D diffusion along DNA to bind other protein and not just DNA targets, producing either *A* + *A* ⇌ *A*_2_ or *A* + *AB* ⇌ *AB* reaction types. Now, excluded volume between proteins or nucleosomes^17^ will impact accessibility for single-file 1D dynamics. Protein binding kinetics in 1D plays an essential role in dwell times^18^ and selectivity for DNA targets^19^, and from the Smoluchowski model for reaction-diffusion dynamics ^20,21^, depends on diffusion constants *D*, intrinsic (barrier) rates for association, *k*_*a*_, and dissociation, *k*_*b*_, and the length *L* of the DNA. Computer simulations of these reaction-diffusion systems commonly use stochastic, particle-based approaches^11,19,22,23^ because random walks in 1D are reentrant, meaning that even in an infinite domain, a diffusing particle will eventually return to its starting point^24^. This renders all standard continuum kinetic descriptions (spatial and nonspatial ^25^) not exact, even for homogeneous systems. Yet these continuum, coarse-grained rate models are widely used because they offer greater computational efficiency^25^. A central question for our work here is therefore not whether these coarse-grained rate models are exact, but when they are accurate enough to be useful and how their effective parameters (*k*_on_ and *k*_off_) should be defined.

Addressing this question requires a quantitative comparison between stochastic particle-resolved dynamics and continuum rate equations, together with a criterion for defining effective rates that is tied to the underlying 1D physics. Here we develop such a framework for reversible binding in 1D. We focus on diffusion-influenced association and dissociation reactions in which molecules move along a filament and can react upon contact, with excluded volume optionally included^26–32^. Using mean-first-passage arguments^26^, we derive a density-dependent effective association rate and define a single-rate approximation evaluated at the initial density. We then compare these coarse-grained descriptions to explicit stochastic reaction-diffusion simulations performed with a 1D implementation of the free-propagator reweighting algorithm^33^ in the NERDSS software package^34,35^. This approach allows us to test both pairwise and many-body systems over a broad range of intrinsic rates, diffusion constants, densities, and domain lengths. In Table 1 we compactly summarize these results via a metric *κ* that measures the ratio of the intrinsic rate to a density-dependent diffusion-limited rate. With this metric we empirically delimit regimes of rate-limited, diffusion-influenced, and diffusion-limited association kinetics.

**Table 1.** *κ* delimits regimes of dynamics from rate-limited to diffusion-limited. Deviations are reported for the ‘worst-case’ scenario of *A* + *A* → *A*_2_ irreversibly, *k*_off_ = 0. *ρ*_*max*_=0.1/nm; max|Δ*A*(*t*)| Eq. 21 is slightly lower at lower densities. Rate change *δ*(*τ*_1/2_) in Eq. 25.

| | Bounds on reaction parameters | $\delta(\tau_{1/2})$ | $\max\vert \Delta A(t)\vert $ | Single rate? |
| --- | --- | --- | --- | --- |
| Rate limited | $\kappa < 0.01$ | $< 13\%$ | $< 3\%$ | $k_{\text{on}}^{\rho_{\text{max}}}$ |
| Weakly diffusion influenced | $0.01 < \kappa < 0.1$ | $13 - 35\%$ | $< 8\%$ | $k_{\text{on}}^{\rho_{\text{max}}}$ |
| Diffusion influenced | $0.1 < \kappa < 1$ | $35\% - 72\%$ | $8 - 14\%$ | $k_{\text{on}}^{\rho_{\text{max}}}$ |
| Strongly diffusion influenced | $1 < \kappa$ | $> 72\%$ | $> 14\%$ | $k_{\text{on}}^{\rho}(\rho)$ |
| Diffusion limited | $10 < \kappa$ | $> 95\%$ | $\sim 17\%$ | $k_{\text{on}}^{\rho}(\rho)$ |

A key finding is that ordinary rate equations (ORE) with a single effective rate can reproduce many-body kinetics surprisingly well in the rate-limited regime. This is despite that for spatial dimensions *d* ≤ 2, a single macroscopic rate constant *k*_on_ does not exist because of reentrant diffusion, as the separation between particles determines association timescales^30^. In *d* = 3 in contrast, the association rate asymptotes to a constant value *k*_on_. In 1D, differences in relaxation kinetics between ORE and reaction-diffusion models of particles emerge far from equilibrium and as equilibrium approaches. Homogeneous but randomly distributed particles initially react very quickly when adjacent, generating spatial correlations that produce slower power-law time evolution for self: *A* + *A* ^29^ and distinct: *A* + *B* ^28^ populations that bind irreversibly (with no volume exclusion), in disagreement with ORE (Fig 1). Differences also emerge for reversible systems in all *d* as they approach equilibrium (obeying Onsager’s regression hypothesis), where diffusion slows the relaxation relative to the exponential relaxation of ORE ^27,36^. However, these differences are all maximized at the diffusion-limit (*k*_*a*_ → ∞) in all *d*, and they shrink as reactions become more rate-limited ^37^. In *d* = 3, the ORE with *k*_on_ becomes an excellent approximation to many-body relaxation beyond the shortest timescales even quite close to the diffusion limit ^30^. In *d* = 2, the regimes of agreement are narrower as defined by the metric 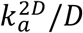^37^ As this dimensionless metric drops below 0.5, the reaction kinetics are only weakly diffusion influenced, and below 0.05 the relaxation is accurately approximated by the ORE with a single *k*_on_^37^. With the loss of another dimension in *d* = 1, we expect even higher sensitivity to spatial correlations, but nonetheless we retain the same emergence of a rate-limited regime.

**Figure 1.**
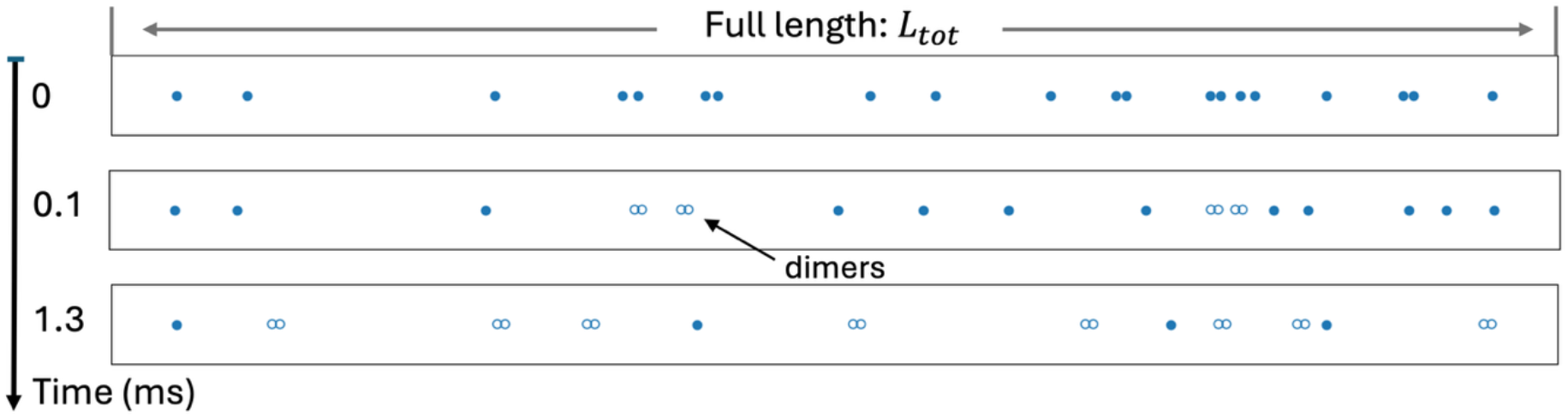
The effective rate ‘constant’ of association in 1D slows down as time progresses and pairwise distance grows. For *A* + *A* → *A*_2_, we illustrate how for an initially homogeneous system of discrete particles at time 0, the reactant monomers (solid circles) that are initially adjacent will rapidly form dimers (open circle pairs). This leads to larger separations as time progresses. While ordinary rate equations predict slower dimerization as the concentration of monomers depletes, in 1D there is an additional effect: the rate ‘constant’ of association is slower as the distance between pairs grows, thanks to reentrant diffusion. Monomers exclude volume with one another, reducing the total accessible volume. If dimers are not annihilated, they will also act as roadblocks that can further impact kinetics.

Another role for simulations here is to assess the role of volume excluding particles, which is typically neglected (although not always^38,39^) by analytical theory that treats particles as points in a dilute limit. In 3D this assumption is broadly valid; the impact of explicit hard sphere particles on their diffusion-influenced reaction kinetics is minimal^40^, where even in the diffusion limit, excluded volume only modestly accelerates association kinetics^25^, or decelerates them for inert ‘crowders’^41^. However, in 1D, volume exclusion is fundamentally distinct and can significantly shift not only kinetics^39,42^ and active phenomena^43^, but the steady-state equilibrium. Instead of simply crowding space, monomers or dimers move in single file and can completely block interactions depending on their linear arrangement along the filament^44^. Excluded volume reduces the explorable space to an effective length *L*_eff_ = *L*_tot_ − ∑*σ*_*i*_ ^45,46^. These effects are generally negligible for diffusive species in 3D or 2D, but not 1D. We measure a quantitative shift in the concentration due to a smaller *L* that thereby shifts the equilibrium to a higher dimer population. Our simulations below further illustrate how the dimers, rather than being annihilated, will also act as physical barriers separating free monomers, which can either slow the relaxation to equilibrium for *A* + *A*, or permanently partition partners relative to their initial linear arrangement in *A* + *B* system. Biologically, returning to 3D via dissociation from the 1D filament thus becomes essential to allow partners to bypass and mix with one another, including in the presence of nucleosome barriers^17^.

In this paper, we first provide relevant background on the theory of diffusion-influenced reactions in *d* = 1, as captured by the Smoluchowski model with reaction collisions ^20,21^. We use mean-first passage time measurements of 1D reaction association events to approximate adaptive, density-dependent rates 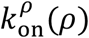 and then a single macroscopic rate constant 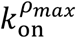 for use in ORE and other continuum models. We define a relatively intuitive metric *κ* that provides empirical and quantitative guidelines in Table 1 for mapping stochastic 1D spatial encounters to efficient, coarse-grained kinetic models that use the effective rate constant. We validate our 1D RD implementation of the FPR algorithm using both pairwise and many-body theory. Our method reproduces known scaling behavior, and illustrates when assumptions about ‘point particles’, or no excluded volume, violate behavior.

Our simulations of reversible dimerization demonstrate that the modified rate equations (MRE)^47^, which use a time-evolving rate from either the Smoluchowski model, 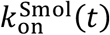, or our density-dependent rate, 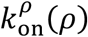, can provide a reasonably accurate description of the spatial relaxation dynamics for homogeneous systems, although both are based on mean-field approximations. Using these simulations and theory allows us to establish quantitative regimes of *κ* where a single rate can be an effective coarse-grained descriptor, depending on tolerated error levels. Through this approach, we provide actionable recommendations for model selection: the MRE with the density-dependent rate 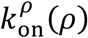 offers a relatively simple yet reasonable approximation for on-the-fly rate evaluation across all *κ* regimes, whereas the single value 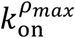 works quite well for small *κ* < 0.01. Collectively, these results demonstrate how despite the intrinsically nonlocal kinetics in 1D, within specific parameter regimes, these 1D reaction dynamics can be simplified into standard biochemical rate equations.

## II. THEORY

### A. Background on Smoluchowski theory for diffusion-influenced reactions

Consider an isolated pair of diffusing particles 1 and 2 in 1D. The separation between them is defined as *x* ≡ |*x*_2_ − *x*_1_|. Reactions follow the Smoluchowski model with the Collins-Kimball radiation boundary condition, and occur upon collision at a binding radius *σ* with an intrinsic/microscopic rate constant *k*_*a*_ ^21^. The limit *k*_*a*_ → ∞ represents the absorbing diffusion-limited reaction boundary. The net diffusion controlling the particle separation *x* is *D*_tot_ = *D*_1_ + *D*_2_, where *D*_1_ and *D*_2_ are the translational diffusion constants of particle 1 and 2, respectively. The probability density *p*(*x, t*|*x*_0_) of finding the particles separated by *x* at time *t* given an initial separation of *x*_0_ is the Green’s function solution to a 1D diffusion equation (1a). With radiation boundary conditions (1c) we have:

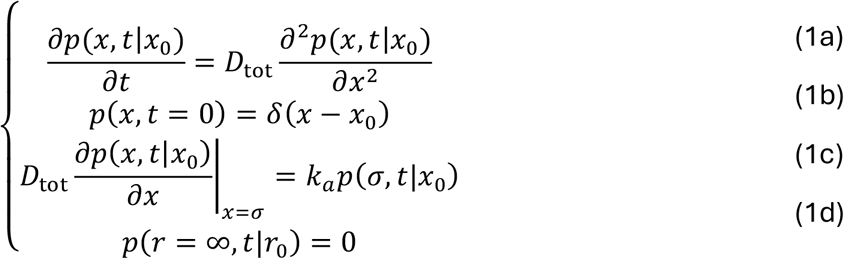

Eq. (1b) defines the delta function initial condition. The radiation boundary condition Eq. (1c) introduces the reaction: when particles collide at *x* = *σ*, they can ‘radiate’ into a bound state with a rate *k*_*a*_, which has unit of nm/μs in 1D. The other boundary condition, Eq. (1d), ensures a normalized probability distribution. Eq. (1) can be solved analytically ^48^, where we refer to it as *p*_*irr*_(*x, t*|*x*_0_) because it captures irreversible association:

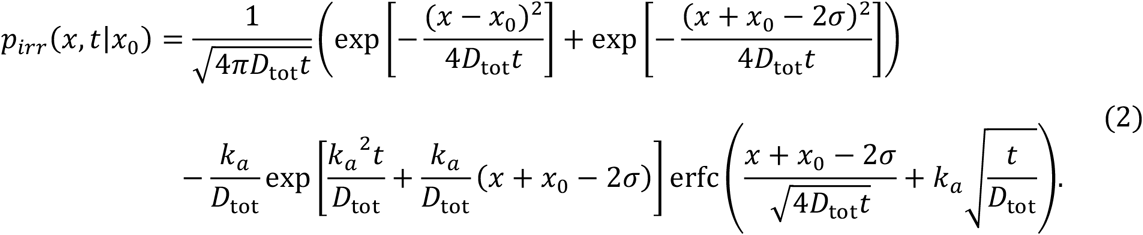

At time *t*, the probability that the reaction has not happened is the survival probability,

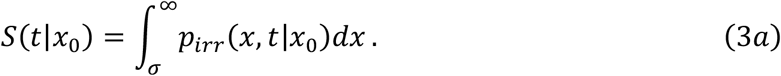

with a solution in 1D given by:

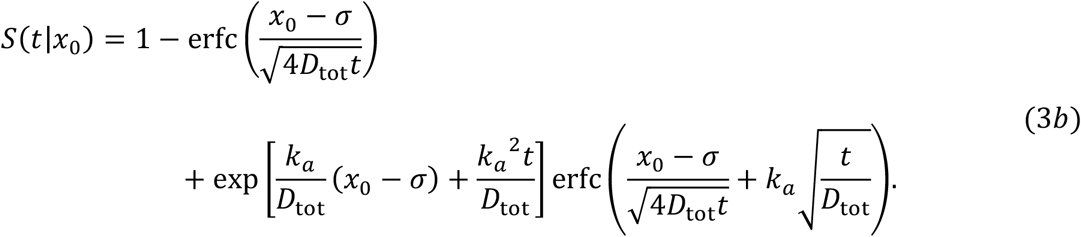

For the fully absorbing boundary condition, where *k*_*a*_ → ∞, the survival probability simplifies to 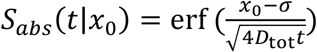. The reaction probability is given by:

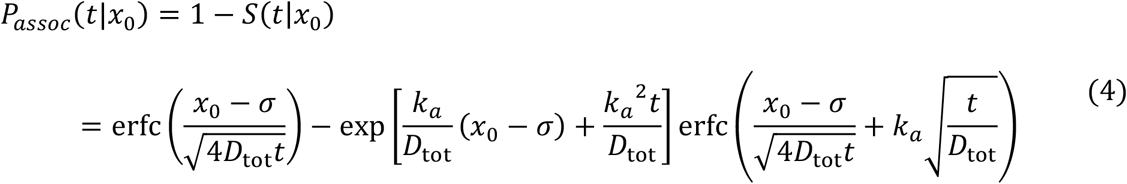

As *t* → ∞, *S* → 0, and *P*_assoc_ → ∞, reflecting the reentrant nature of diffusion in *d* = 1; despite the infinite domain (Eq. 1d), particles will always end in the bound state.

The survival probability of Eq. (3) can be used to define a *macroscopic*, time-dependent rate of association based on an initial state of unbound reactants at contact:

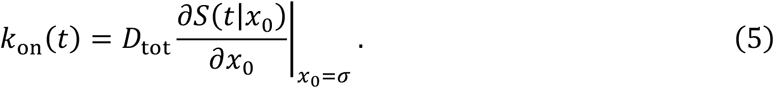

For radiational boundary condition (Eq. 1c), Eq. 14 can be also rewritten as *k*_on_(*t*) = *k*_*a*_*S*(*t*|*x*_0_ = *σ*) ^21,49^. In 1D, this rate was first derived by Collins and Kimball to give ^21^:

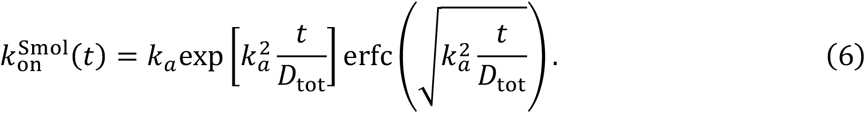

For reversible systems, we have the well-known equilibrium association constant is given by:

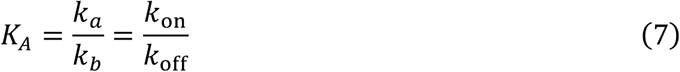

where *k*_*b*_ is the intrinsic or microscopic rate of dissociation back to contact at *σ*. For time-dependent macroscopic rates *k*_on_(*t*), we therefore must use a time-dependent dissociation rate defined using Eq. 7, where the intrinsic (barrier) rates *k*_*a*_, *k*_*b*_ are always constant.

At the diffusion limit (*k*_*a*_ = ∞), Eq. 6 reduces to the rate for the absorbing BC of the original Smoluchowski model, 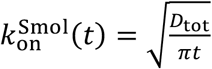. In 1D, we also see that the macroscopic rate asymptotes to this same functional form at long times for all *k*_*a*_ values:

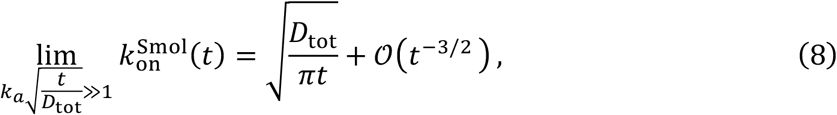

At short times, 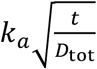 is small and we have:

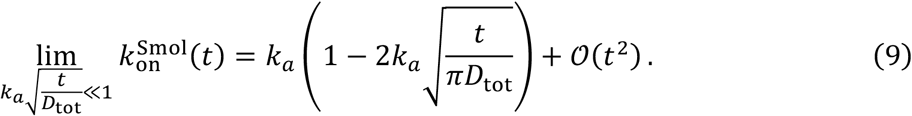

### B. Excluded volume of particles reduces the domain size nonnegligibly in 1D

The many-body theory and simulations are restricted to a 1D domain length of *L*_tot_. Although the many-body theoretical results we use below always assume point particles, in simulations, all reactive pairs exclude volume at a separation of |*x*| < *σ*. These hard-core exclusions can also be extended to the bound dimer states (*A*_2_ or *AB*). For annihilation reactions, the bound dimer states will not exclude volume, but for the reversible systems, we will enforce excluded volume by the bound states. However, even when we make bound states invisible, excluded volume between monomers can noticeably shift the kinetics and equilibrium, even for density fractions well below *σN*/*L*_tot_ = 0.5, with *N* the total number of monomers. Generally, the accessible domain length *L* is therefore dependent on the reactant type, total partners present, and their relative *σ*, as the linear arrangement of particles impacts whether they can ‘pass through’ one another. For each reactant type, its accessible domain length *L*_*r*_ is reduced relative to the full defined domain via the number *N*_ex_ of particles in the system that exclude volume with the reactant *r* with a binding radius *σ*_*i*_:

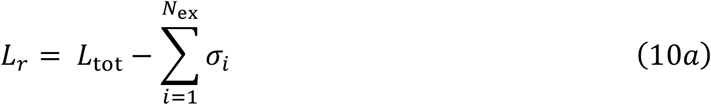

For a homo-dimerization reaction between *A* molecules there are only two species, *A* and *A*_2_ and if all 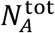 molecules occupy space throughout time this expression simplifies to:

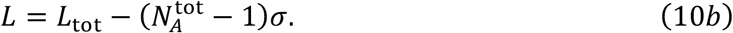

We assume here that the dimer state should continue to exclude volume with all other species. In our simulations, the monomer centers are placed at the binding radius separation *σ* when in the dimer state, so that upon dissociation, they are already excluding volume. Therefore, if dimers continue to exclude volume, then all 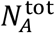 molecules occupy space through the entire simulation. For irreversible homodimerization with annihilation, the accessible domain length will return to *L*_tot_ as a function of time as monomers are removed from the system (*N*_*A*_(*t* → ∞) → 0), with *L*(*t*) = *L*_tot_ − (*N*_*A*_(*t*) − 1)*σ*.

### C. Derivation of a density-dependent rate constant to approximate 1D reversible binding

Theoretically, the rate of association between a pair of reactants in 1D is always sensitive to their separation, and no single rate constant exists. The mean-field Smoluchowski model predicts a time-dependent rate in section A (Eq. 5–6), but this is based on assertion of a fixed initial separation *σ* and subsequent diffusion in an infinite domain. We here define a density-dependent rate as an alternate approximation for estimating a time-evolving rate of association. This rate model assumes a homogeneous distribution of particles, but allows for changes to the relative separation between them. We follow the approach we used in 2D ^37^, by starting with the mean-first passage time (MFPT) previously derived by Szabo and co-workers for a single reactant pair in a bounded domain ^26^. A stationary particle is placed at *x* = 0 with radius *σ*, and a single partner *B* with diffusion constant *D* searches within the domain bounded by a reflective boundary at *x* = *L*_*tot*_, giving an accessible domain size of *L*_*tot*_ − *σ*. By averaging over all possible initial positions for *B* uniformly in this domain, the MFPT is given by: 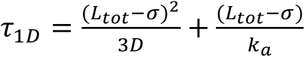^26^. We can then define the mean macroscopic rate via 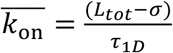. This mean macroscopic rate therefore averages over fast events for *B* initially adjacent to its partner, and the slow events for *B* at the maximal separation *L*_*tot*_ − *σ*, and it accounts for diffusion within a bounded domain.

However, for a many-body system, the average separation between each single reactant pair is not defined by the full domain length but instead is dependent on the density *ρ* of the reactants. We therefore define the mean density *ρ*(*t*) = max(*N*_*A*_(*t*), *N*_*B*_(*t*))/*L* for a reaction *A* + *B* ⇌ *AB*, where *N*_*A*_(*t*), *N*_*B*_(*t*) are the copy numbers of molecules *A, B* at time *t*, respectively. For a homodimer reaction, *ρ*(*t*) = *N*_*A*_(*t*)/*L*. The domain length here can represent the accessible domain length defined in Eq. 10 based on volume exclusion. We can then define a density-dependent rate using the MFPT within this shorter lengthscale, which can therefore be a function of time, as:

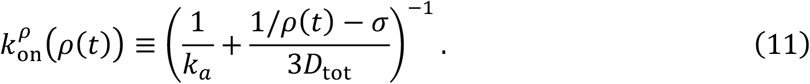

with *D*_tot_ = *D*_*A*_ + *D*_*B*_. This rate should be interpreted as predicting the mean association rate assuming particles are homogeneously distributed at a density *ρ*(*t*) and therefore constrained by a lengthscale *l* = 1/*ρ*. It therefore does not describe the fastest events or slowest events, but rather the average speed for this domain length. Importantly, for reversible reactions, the dissociation rate is also density dependent, 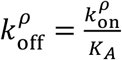.

To simplify further, we can define a single macroscopic rate by choosing a specific particle density. We propose the density when the system is most densely populated, at the initial time, although for steady-state systems, one can use the steady-state density. By choosing the maximal density, we will show below that the half-time of the spatial relaxation kinetics is well approximated, as this mean macroscopic rate does not capture the fastest early events (short-time relaxation), or the slowest late events (long-time relaxation), but reaches 50% completion at similar timescales for irreversible systems. Using the density *ρ*_max_ = max(*N*_*A*,tot_, *N*_*B*,tot_) /*L* to define an average domain size produces the single rate constant:

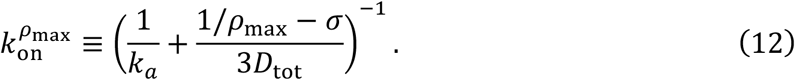

The functional form for Eq. 11 and 12 closely mirror the truly steady-state rate one derives in 3D: 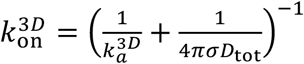. In 3D one adds together the times of two processes, diffusing to contact with the well-known rate *κ*_*D*_ = 4*πσD*_tot_, and crossing the barrier once at contact with rate 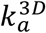, to define the net macroscopic rate. By analogy, we can see in 1D that the purely diffusion-limited rate to contact is given by:

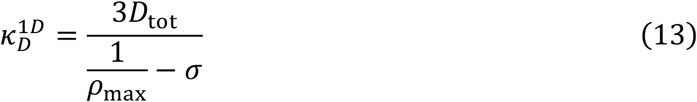

albeit here it is defined by necessity for a bounded domain, unlike in 3D. We show below how this approximation for a single macroscopic rate, when used in ordinary rate equations, compares to the full single-particle reaction-diffusion kinetics.

### D. Metric *κ* for assessing regimes of rate vs diffusion-limited association

In general, one expects that the smaller the value of *k*_*a*_, the more rate-limited the association kinetics will be in any dimension. For increasingly rate-limited association, the time evolution of the reaction dynamics becomes increasingly independent of diffusion and domain size, and more accurately approximated by a single macroscopic rate. We define a relatively intuitive metric to establish regimes of rate-limited vs diffusion-influence kinetics in 1D by taking the ratio of the intrinsic barrier rate *k*_*a*_ with the diffusion-limited 1D rate defined by Eq. 13 or equivalently Eq. 11 when *k*_*a*_ → ∞ to get:

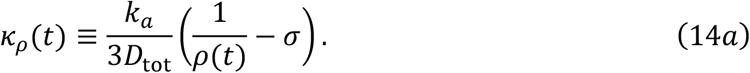

While this value thus varies with time/system density, for a given set of parameters, we will refer to the single *κ* value as the value initially at the maximum density *ρ*_max_, similar to our definition of the macroscopic rate, to get:

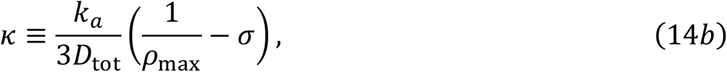

For reversible reactions, *κ*_eq_ = *κ*_*ρ*_(*t* → ∞) will be the largest value reached for a system initialized with all molecules as monomers, such that *κ* ≤ *κ*_*ρ*_(*t*) ≤ *κ*_eq_. With *κ* ≫ 1 the system is diffusion-limited, and *κ* ≪ 1 is rate-limited. With *κ*_eq_ ≪ 1 this ensures the dynamics is always rate limited even at the slowest relaxation regimes. In Table 1 we are more specific than just *κ* ≪ 1 based on our analysis in the results.

### E. Reactants approaching from two sides requires doubling the association rate

Both the single-pair RD model (Eq. 1) and the (approximate) many-body Smoluchowski model (Eq. 6), assume that a reactant *A* can only be bound by partners *B* approaching from one side, e.g. [*σ*, +∞). More generally, however, multiple reactive pairs can exist within the domain *L*, producing reactive flux across both sides of the reactant molecule A, from the left and the right. If the reactants are distinct (*A* + *B* systems), comparison of the single-particle reaction-diffusion kinetics parameterized by *k*_*a*_, *D*, and *σ* to nonspatial rate equations requires the macroscopic rate *k*_on_ calculated using Eq.6, Eq. 11, or Eq. 12 must be multiplied by a factor of 2 in ORE or MRE. In contrast, this factor of 2 is not needed for self-binding *A* + *A* systems. This also changes the equilibrium constant exclusively for these two-sided *A* + *B* systems, giving 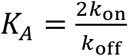.

## III. METHODS

### A. Numerical method to solve the 1D many-body problem uses the FPR algorithm

#### A1. FPR for 1D pairs

To numerically solve the 1D reaction-diffusion many-body problem (Fig 1), we break the full system into the constituent pairwise interactions that obey the tractable 2-body problem defined in Eq. (1). We here use the Free-Propagator Reweighting (FPR) algorithm, which was derived and described in detail in ref ^33^. Here we briefly explain the necessary function evaluations and their definitions in 1D. FPR reproduces the exact reaction probability, *P*_*assoc*_(*t*|*x*_0_) for each particle pair, and rigorously obeys detailed balance for reversible reactions. The algorithm samples position updates using the free propagator while preserving excluded volume (*x* ≥ *σ*) but uses trajectory reweighting with the correct propagator *p*_*irr*_(*x, t*|*x*_0_) to recover the correct *P*_*assoc*_(*t*|*x*_0_) defined in Eq. (4). The reweighting factor *w*_*ratio*_ is given by:

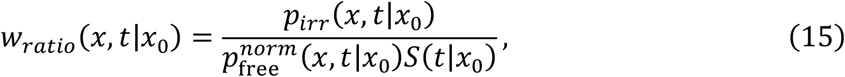

where 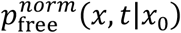 is the solution to the diffusion equation without a reactive boundary (Eq. 1a, 1b, and 1d only) but normalized after excluding all separations where *x* < *σ*. Numerically this is implemented by rejecting all position updates with *x* < *σ* and resampling until *x* ≥ *σ*. This solution is given by:

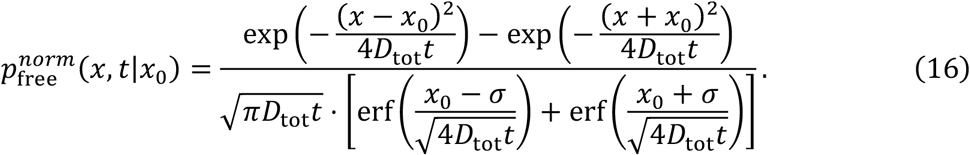

In Eq. (15), *p*_*irr*_(*x, t*|*x*_0_)/*S*(*t*|*x*_0_) is the normalized distribution of a pair of molecules obeying Eq. 1. For each reactive pair, we therefore evaluate reaction probabilities at each step using the reweighted association probability given by:

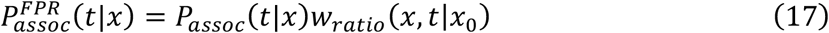

where *t* = *N*Δ*t* is the time after *N* steps of length Δ*t*. To apply trajectory reweighting, the reweighting factor *w*_*ratio*_ is accumulated throughout the entire trajectory by exploiting the Markovian property of diffusional dynamics, starting from when the particles first enter the reaction zone (defined below) at *x*_0_ ≤ *R*_*max*_. ^33^, namely,

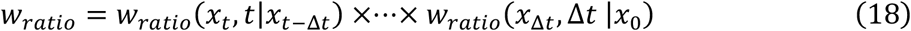

We initialize *w*_*ratio*_ = 1 when the particles first enter the reaction zone.

Because FPR involves trajectory reweighting, it can suffer from under-sampling errors if free trajectories are not sampled, and therefore not reweighted, in regimes where the reactive trajectory is nonnegligible, as detailed previously ^33,37^. This trajectory overlap is primarily an issue for very long reactive trajectories. However, because reactions only occur upon collision, the reaction probabilities *P*_*assoc*_(*t*|*x*) drop to zero over relatively short distances, meaning trajectories revert to free diffusion relatively quickly. Once *P*_*assoc*_(*t*|*x*) drops to zero at a separation *x* = *R*_*max*_, the particle pair are no longer evaluated for reactions, as they are simply free diffusing, and reweighting can be reset (*w*_*ratio*_ = 1). We choose the value of *R*_*max*_ based on the average separation between a freely diffusing pair scaled by a heuristic prefactor *c* that ensures the association probability is low enough to be neglected:

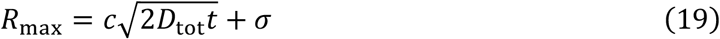

where in 1D we set *c* = 4.

To capture dissociation occurring with the intrinsic rate *k*_*b*_ (which does not include diffusion apart), we model this process as Poissonian, producing a probability:

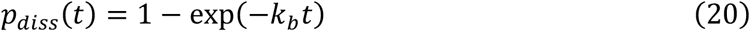

To retain detailed balance, a dissociation event must return the particles back to a separation *x* = *σ*, as association reactions only occur at this collision distance. This ensures that at equilibrium, whether in 3D, 2D or 1D, the relation holds in Eq. 7.

We note a technical detail for recovering macroscopic rates of self-binding (*A* + *A*) reactants. In both 3D and 2D, to recover the expected macroscopic rate of self-binding reactants and enforce that *K*_*A*_ = *k*_*a*_/*k*_*b*_, during the Green’s function evaluation of the reaction probability, NERDSS internally replaces *k*_*a*_ → 2*k*_*a*_. In 1D, however, because the reactants access partners from two sides, the flux is naturally doubled and in NERDSS we evaluate Eq. 4 with simply *k*_*a*_.

#### A2. Implementation details for many-body 1D RD in NERDSS

Our 1D reaction-diffusion simulator is implemented in NERDSS and follows the same reaction and propagation steps for many-body systems in 3D, 2D, or transitioning between ^35^. The software is open source at https://github.com/JohnsonBiophysicsLab/NERDSS. Briefly, in each fixed time step Δ*t*, we first evaluate dissociation reactions for any bound pair using Eq. 20. We then identify all reactive molecule pairs where *x* < *R*_*max*_. We evaluate the reaction probability for each of these pairs using Eq. 17. For each particle we perform maximally one reaction of any type, by comparing reaction probabilities to a URN. For molecules that have not reacted in each Δ*t*, we propagate them by Brownian motion with rejection and resampling whenever *x* < *σ*. When monomers form dimers, they are always placed such that their centers are exactly separated by *σ*. Thus, upon dissociation, the now-free monomers are excluding volume with one another.

In NERDSS, we currently define 1D filaments as straight lines parallel to the x-axis spanning from one end to the other end of the water box. A molecule is assigned to moving on these filaments with the tag “isPromoter=true” and therefore undergoes 1D reactions with similarly tagged particles. For 1D molecules, NERDSS only propagate their *x* coordinates according to the *x* component of the diffusion constant, *D*_*x*_, and uses *D*_*x*_ to determine the reaction probabilities. For interacting molecules, we enforce excluded volume and ‘hopping over’ events between reactive pairs by canceling all propagations that causes 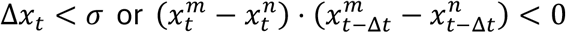 between molecules *m* and *n*. If two molecules do not interact, by default they do not exclude volume and ‘pass through’ one another. To enforce excluded volume for all pairs, we typically introduce additional interfaces to each monomer that react with a rate *k*_*a*_ = 0. By default, all 1D molecules are initialized at the center of the water box, i.e. *y* = 0 and *z* = 0, with random *x* coordinates. They can be assigned at different positions by using the “-c” or “--coordinate” flag when running simulations. To define different filaments at different positions, one can assign different *y* and *z* values to 1D molecules in “fixCoordinates.pdb”.

#### A3. Choosing the time-step Δt

In FPR, the fixed time step Δ*t* is chosen to best recapitulate isolated 2-body problems based on the total reactive particle density *ρ*, such that each particle does not ‘see’ multiple particles at once. At this Δ*t*, each reactive particle should have, on average, at most 1 partner within *R*_max_. We therefore set out to solve the equation *N*_*rct*_ ≥ *ρ*(2(*R*_max_ − *σ*)) with *N*_*rct*_ = 1. The factor of 2 is necessary in 1D because in general, reactant partners can access binding reactions from both the right and the left of the particle, creating two interfaces for reactive flux. Considering the most dense reactant specie *N* as limiting, we have *ρ* = *N*_max (*A,B*)_ /*L*, and we find that 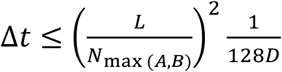. If the timestep is larger than this, simulations can fail to reproduce the correct kinetics or equilibrium.

### B. NERDSS simulation details

#### B1. Simulations of 1D reactive pairs

Two molecules are initialized at *x*_1_ = −1 nm and *x*_2_ = *x*_1_ + x_0_ on a filament with *L*_tot_ = 500 nm to mimic an infinite system. We fix *σ* = 1 nm, *Δt* = 0.1 *μ*s, monomers diffuse with constant *D* = 0.1 nm^2^/*μ*s, and *h*^2^ = 1 nm^2^. We vary x_0_ and *k*_*a*_ in different experiments as shown in Fig. 2. In each experiment, *N*_rep_ = 10000 trajectories are repeated. For one trajectory, the simulation is terminated when association happens or the particles exit the reaction zone, i.e Δ*x* > *R*_max_ = 1.8 nm. The values of *w*_*ratio*_ and Δ*x* at the step before exiting the reaction zone is recorded in wratios.csv for analysis.

**Figure 2.**
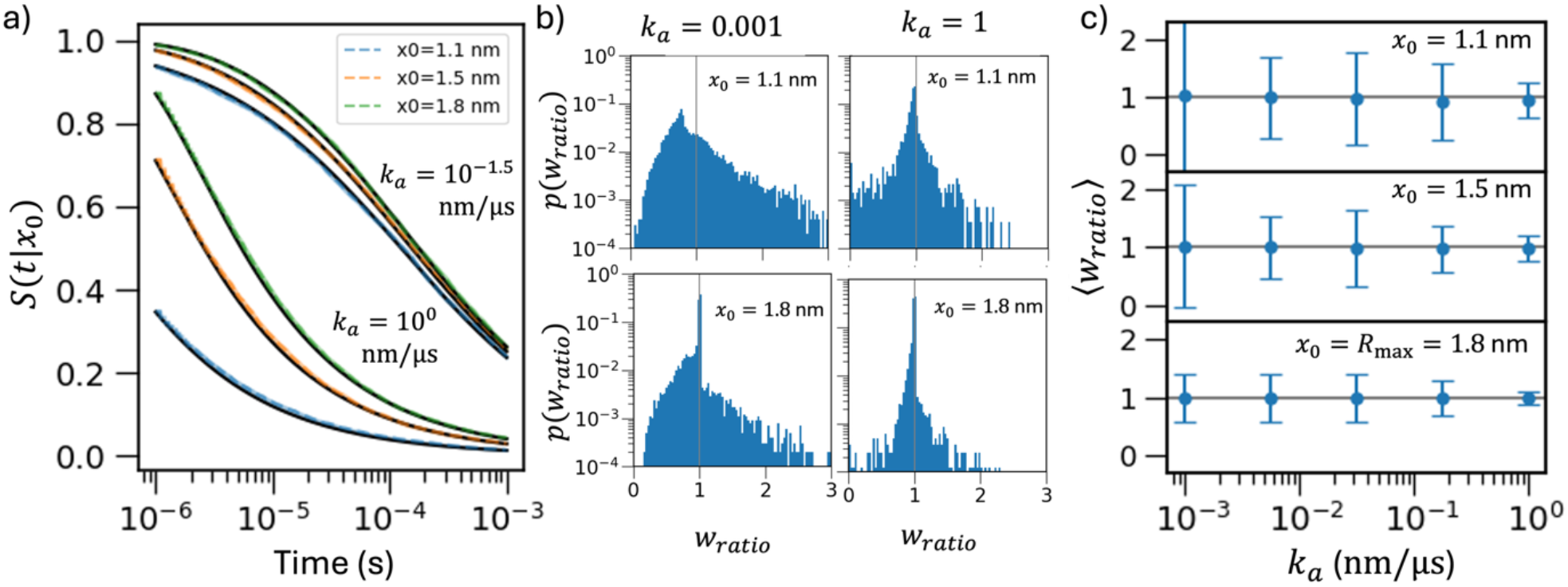
NERDSS 1D simulations of reactive pairs accurately reproduces the theoretical survival probabilities over time. Simulations of two particles are performed with time step *Δt* = 0.1 *μs*, contact radius *σ* = 1 *nm*, and each molecule has *D* = 0.1 *nm*^2^/*μs*, giving *D*_*tot*_ = 2*D* = 0.2 *nm*^2^/*μs* and *R*_*max*_ = 1.8 *nm* (Eq. (8)). a) For both *k*_*a*_ = 0.032 (upper curves) and 1 *nm*/*μs* (lower curves), we find excellent agreement between the survival probability from NERDSS simulations (colors) and the theory (black lines-Eq. 3). We validated for several initial starting separations *x*_0_ = 1.1 (blue), 1.5 (orange), and 1.8 nm (green). We performed 7200 trajectories of single pairs. The standard errors (shown—below line width) are calculated with 500 times bootstrap resampling. b) We verified that the distributions of reweighting ratios *w*_*ratio*_ calculated as each pair exits the reaction zone at *R*_*max*_ are centered around 1 and are not too broad. Results for *k*_*a*_ = 0.001 *nm*/*μs* in left column, and for 0.1 *nm*/*μs* in the right column. C) We report the ⟨*w*_*ratio*_⟩ across all trajectories started from the same *x*_0_. The means are all very close to 1, as expected. Error bars report the standard deviation, with larger variance for smaller *x*_0_.

#### B2. Simulations of many-body irreversible association

***A*** + ***A*** → ∅ In simulations, 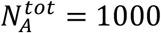 molecules are initialized randomly on a filament with length *L*_*tot*_ = 2 *μ*m, 5 *μ*m, or 10 *μ*m. We fix *σ* = 1 nm, *Δt* = 0.1 *μ*s, and monomer diffusion constant *D* = 0.1 nm^2^/*μ*s (so *D*_tot_ = 0.2 nm^2^/*μ*s). Now, with *L*_*tot*_ ≥ 2 *μ*m, *k*_*a*_ = 10 nm/*μ*s is selected to push associations to the diffusion limit (Table 1). The number of time steps is 2 × 10^6^ to achieve a wall time at 0.2 s. For each system we collect statistics over 48 repeated trajectories and plot averages along with the SEM at each time point, 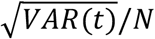.

#### B3. Simulations of reversible association

***A*** + ***A*** ⇌ ***A***_**2**_ In simulations, 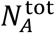 molecules are initialized randomly on a filament with length *L*_tot_ in the monomeric state. We set the diffusion constant of A molecules to be *D* = 0.1 nm^2^/*μ*s (so *D*_tot_ = 0.2 nm^2^/*μ*s) and *σ* = 1 nm. The number of time steps is selected to target a wall time at 25 s to reach equilibrium. The A species excludes volume in both the monomer and dimer (bound) state. For each system we collect statistics over 48 repeated trajectories.

#### B4. Simulations of many-body irreversible association

***A*** + ***B*** → ∅. In simulations, an equal number of *A* and *B* molecules are initialized randomly on a filament. The total number of *A* molecules is 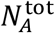. We fix *σ* = 1 nm, *Δt* = 0.5 *μ*s, and monomer diffusion constant *D* = 0.2 nm^2^/*μ*s (so *D*_tot_ = 0.4 nm^2^/*μ*s). By definition, the dimer is annihilated and does not exclude volume. Furthermore, A does not exclude volume with other A molecules, and likewise for B to B. Only monomeric A and B exclude volume with one another. Now, with *L*_*tot*_ ≥ 2 *μ*m and ensuring 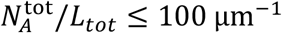, *k*_*a*_ = 100 nm/*μ*s is selected to push to the diffusion limit. The number of time steps is 2 × 10^8^ to achieve a wall time at 10 s. Statistics for each system are averaged over 48 trajectories.

#### B5. Simulations of reversible binding to a target at the end

***A*** + ***B*** ⇌ ***AB*** Each simulation contains 9 filaments and is repeated by 96 trajectories. Each filament has 1 fixed A molecule (the target) initialized at the left end. 20 B molecules are moving along the whole filament. The initial coordinates of B molecules on 9 filaments for 96 trajectories are generated randomly by a uniform distribution without overlap. The simulations read in 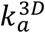, and convert to the 1D value using: 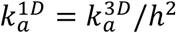, with *h*^2^ = 1 nm^2^. *D*_*A*_ = 0, *D*_*B*_ = 0.1 *μm*^2^/*s*. *K*_*A*_ = *k*_*a*_/*k*_*b*_. Rate equations solve: 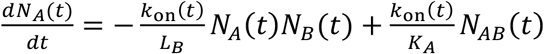. Here we consider not only the excluded volumes between *A* and *B* molecules, but between *B* molecules in the unbound and bound states. The accessible domain length for sampling of the *B* molecules is therefore reduced to *L*_*B*_ = *L*_*tot*_ − *N*_*B,tot*_*σ*_*AB*_. Another set of simulations allowed the target to diffuse with *D*_*A*_ = *D*_*B*_.

#### B6. Simulations of reversible binding to a target at the center

***A*** + ***B*** ⇌ ***AB*** Each simulation contains 9 filaments and is repeated by 96 trajectories. Each filament has 1 fixed A molecule (the target) initialized at the center. 20 B molecules are placed half on the left of the target and half on the right of the target. B molecules diffuse within their half without crossing over the central target A. The B molecules do *not* exclude volume with one another, they only exclude volume with the single A target. The simulations read in 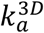, and convert to the 1D value using: 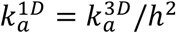, with *h*^2^ = 1 nm^2^. *D* = 0, *D* = 0.1 *μm*^2^/*s*. *K* = 2*k*_*a*_/*k*_*b*_. Rate equations solve: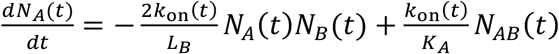. We again consider not only the excluded volumes between *A* and *B* molecules, but between *B* molecules in the unbound and bound states. *L*_*B*_ is the same as in (B5). Another set of simulations allowed the target to diffuse with *D*_*A*_ = *D*_*B*_.

### C. Numerical Methods and Analysis

We solve the MRE numerically using the SciPy package solve_ivp with the ‘LSODA’ integration routines^50^ in Python3, and with ode23s in MATLAB. With constant rates (ORE), we use the analytical solutions. To compare kinetics from two distinct models, the MRE with the Smoluchowski rates and the ORE with a single *k*_on_, we normalize the time-dependent absolute error:

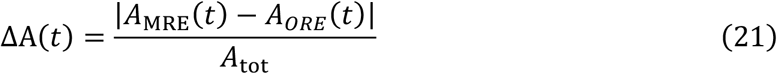

where *A*_tot_ is the total number of *A* molecules at time zero.

## IV. Results

### A. The FPR algorithm accurately captures single-particle reaction-diffusion dynamics for a single pair in 1D

We first establish here that our numerical method (the FPR algorithm in the NERDSS simulation software) for simulating single-particle reaction-diffusion dynamics in 1D accurately reproduces the expected theoretical behavior for simple systems. For a single pair of reactants parameterized by *k*_*a*_, *D*, and *σ* the simulations should recover the theoretical survival probability given by Eq. 3 for any initial separation *x*_0_, which is an exact solution including volume exclusion at *σ*. We find excellent agreement in Fig. 2a, evaluated for *k*_*a*_ = 1 and 0.03 *nm*/*μs*, at several initial separations. Because the FPR algorithm samples from the free propagator rather than the exact Green’s function, trajectory reweighting is necessary to recover the exact association probabilities between all reactive pairs (Eq. 17). The reweighting ratios, *w*_*ratio*_, are accumulated for each pair while they remain in the reaction zone defined by *x* < *R*_*max*_. The distribution of reweighting ratios (Fig. 2b) are expected to average to 1 when the sampled trajectories are fully sampling from the exact theoretical trajectories^33^. As expected, we find ⟨*w*_*ratio*_⟩ is close to 1 for all simulations (Fig. 2c). The width of the distribution of *w*_*ratio*_ values is shown by the error bars in Fig. 2b and tends to increase for smaller *k*_*a*_ values. This is due to the longer trajectories required to finally perform an association reaction as *k*_*a*_ → 0; longer trajectories accumulate more reweighting factors simply by taking more steps, creating more separation between the exact and sampled distributions of trajectories.

### B. NERDSS simulations accurately reproduce many-body reaction dynamics for *A* + *A* → ∅

To validate our numerical implementation of many-body FPR in the NERDSS software, we further performed simulations of the annihilation reaction *A* + *A* → ∅ with periodic boundary conditions ^29^. At the diffusion limit (*k*_*a*_ → ∞), all collisions between A particles lead to a reaction, and the dimers that would normally be created are instead annihilated. An exact analytical solution to this 1D problem assuming a dilute system of point particles^29^ is given by:

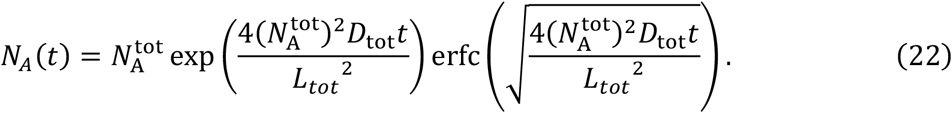

This theoretical result does not capture excluded volume. In our simulations, the monomeric *A* species all exclude volume with one another (although dimers are invisible), which does reduce the accessible domain length below *L*_*tot*_ as noted in Eq. 10. With a sufficiently low volume fraction of molecules at 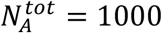 molecules in *L*_*tot*_ = 10 *μ*m, where 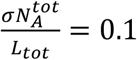, the simulations agree nearly perfectly with the theory in Eq. 22 (Fig 3). As we increase the concentration by reducing *L*_tot_ to 5 *μ*m or 2 *μ*m, 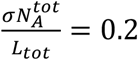 and 0.5, we see the long-term association kinetics is again nearly perfect—the system has substantially diluted by this point--but at shorter times, the simulations are slightly faster than the dilute theory because of their excluded volume.

**Figure 3.**
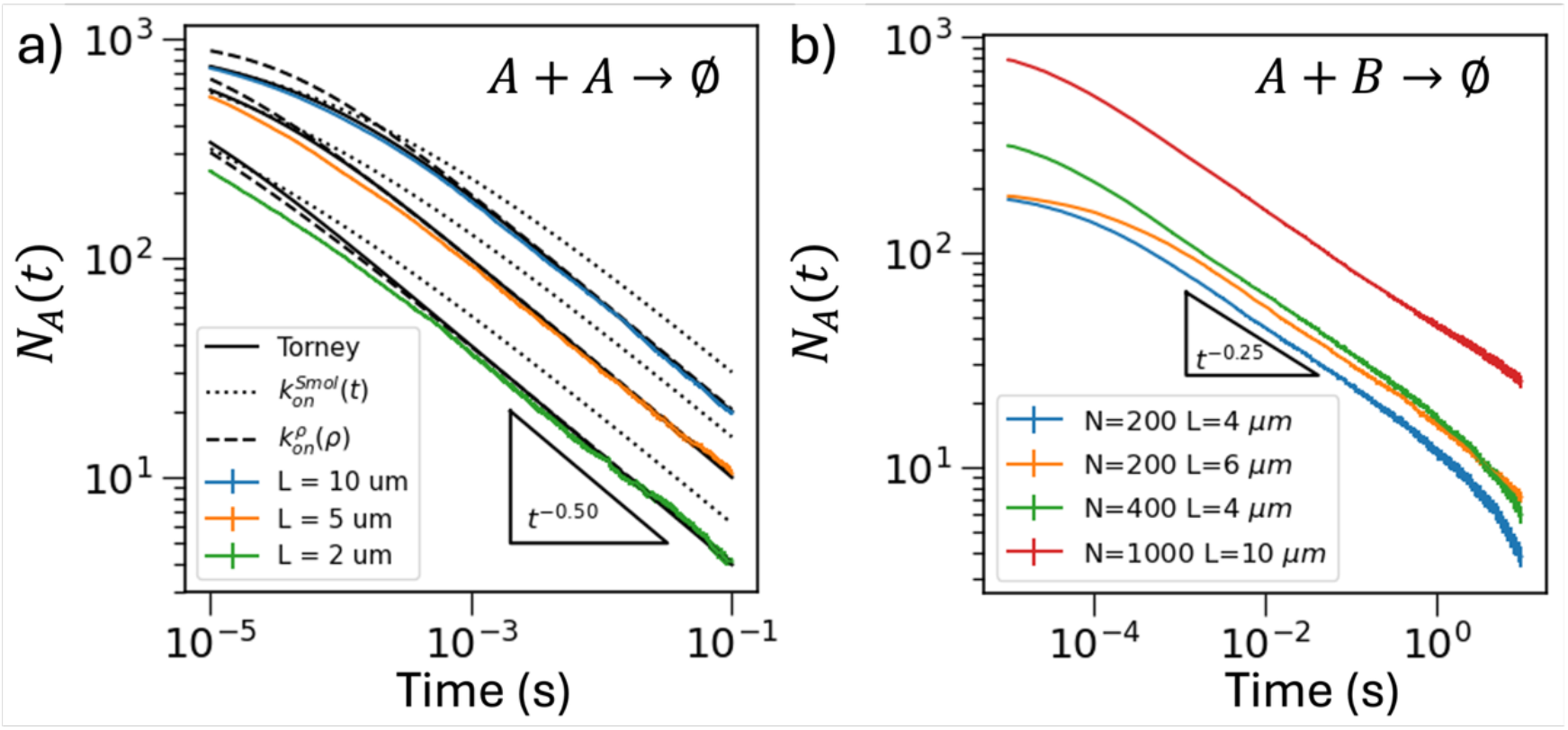
NERDSS simulations of many-body diffusion-limited annihilation dynamics reproduce theoretical predictions. We compare NERDSS simulations (colored curves with error bar reporting the standard error of the mean (SEM(*t*))) with theoretical solutions in the diffusion-limit *(k*_*a*_ → ∞) that account for spatial correlations. a) For *A* + *A* → ∅, NERDSS simulations at 3 densities (blue=0.1/nm, orange=0.2/nm, green=0.5/nm) reproduce the expected power-law decay *N*_*A*_(*t*)~*t*^−1/2^, as illustrated by the black triangle. The exact (dilute) many-body theory of Torney is shown in solid black (Eq. 14). The MRE (Eq. 15) with the time-dependent Smoluchowski rate is in dots, and the MRE with our density-dependent rate is in dashed. Simulation parameters are: *N*_*rep*_ = 48, 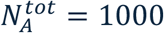, *σ* = 1 nm, *Δt* = 0.1 *μs, D* = 0.2 *nm*^2^/*μs*. We used a finite *k*_*a*_ = 10 *nm*/*μs*, but it well-approximates the diffusion-limit effectively for these densities (Methods). b) For *A* + *B* → ∅, with equal populations of A and B reactants, we compare our many-body simulation with the asymptotic law *N*_*A*_(*t*)~*t*^−1/4^ (the reference triangle). To mimic theoretical assumptions, *A* particles do not ‘see’ other *A* particles, and same for *B* to *B*. Only *A* and *B* exclude volume with one another with *σ* = 1 nm. Green and red curves have the same particle density, but the red curve has a larger *L* and *N*, with red showing a larger regime of power-law decay. Both green and blue have the same domain length *L* (but distinct densities), showing a transition to faster decay at a similar time. Simulation parameters are: *N*_*rep*_ = 48, *Δt* = 0.5 *μs, D*_*A*_ = *D*_*B*_ = 0.1 *nm*^2^/*μs*. We use a large finite *k*_*a*_ = 1000 *nm*/*μs* that evaluates reaction probabilities consistent with the diffusion-limit (Methods). After a time duration of *T* = 10 s, we have the average relative displacement 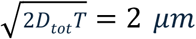, or half the domain length of the blue and orange curves, driving the transition to a faster decay.

### C. Our derived density-dependent rate constants can predict the reaction dynamics for *A* + *A* → ∅ with reasonable accuracy

For these same many-body simulations, we also compared them to the prediction of more approximate theories: the continuum rate equations using two different time-evolving rate models, Smoluchowski’s model, 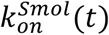 (Eq. 6) and our density-dependent rate model, 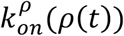 (Eq. 11), by solving the modified rate equation (MRE):

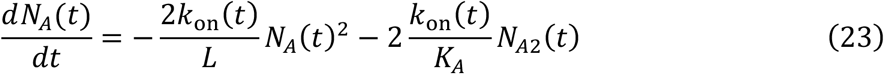

where for irreversible reactions we have 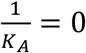. The MRE is not exact due to the mean-field approximations inherent to *k*_on_(*t*), which also ignore excluded volume. For constant rates (*k*_on_(*t*) = *k*), this simplifies to the ORE. As we describe in Theory section II.C, our density-dependent rate, 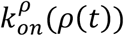 is based on the mean-first passage time for a single diffusing particle to react with a stationary partner in a bounded domain, *L*_2_. For many-body systems, we therefore adapt the mean association rate by redefining the average domain length *L*_2_ between particles as a function of homogeneous reactant density *ρ*(*t*), producing a time-evolving rate, rather than a constant rate. One aim of introducing this new rate model is that, unlike the Smoluchowski model, it does not require a specific set of initial conditions and therefore can be used under more general nonequilibrium conditions where density can both rise and fall. We can capture one element of volume exclusion by using the reduced domain length, *L* from Eq. 10.

As noted previously by Torney et al^29^, solving the MRE with Smoluchowski’s time-resolved rate (at the diffusion limit) underestimates the speed of association somewhat at longer times (Fig 3). This is because the Smoluchowski rate model assumes that particles can diffuse apart to infinite separation (this model has no *L* dependence), and it neglects many-body correlations, whereas here the monomers are in a bounded domain. Nonetheless, this irreversible model does capture the correct form of the *t*^−1/2^ asymptotic behavior consistent with Torney’s solution and the NERDSS simulations.

Solving the MRE using our density-dependent rate 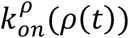 produces excellent agreement to the true RD kinetics at long times for this irreversible reaction (Fig 3). Initially, the density-dependent rate predicts slower association relative to the simulations, as expected by its derivation. This is because the beginning of the simulations is precisely when adjacent pairs drive the fastest possible association events that exceed the mean association rate (Fig 1). As the system relaxes, the good agreement of the MRE to the simulations indicates that using the time-dependent mean density to predict the association rate can predict collision times between pairs. Although the density-dependent rate method assumes reflective boundary and Torney’s theory assumes periodic boundary, in the case of self-association, the difference between their asymptotic dynamics is minimal (Fig. 3a). This time-evolving rate model also captures the expected *t*^−1/2^ asymptotic behavior^30^, consistent with Torney’s solution and the NERDSS simulations.

### D. NERDSS simulations accurately reproduce many-body reaction dynamics for *A* + ***B*** → ∅

For the heterodimerization annihilation reaction, *A* + *B* → ∅, there is no simple closed-form theoretical solution for the full time-dependent RD dynamics. However, for diffusion-limited reactions (*k*_*a*_ → ∞), analytical theory in the dilute (point particle) limit predicts that with an equal amount of *A* and *B* reactants homogeneously distributed, the long-time asymptotic relaxation on an infinite filament is proportional to *t*^−*d*/4^, not *t*^−*d*/2^ of the previous section^28^. It does not predict short-time relaxation. Here we validated the long-time relaxation with our simulations, reproducing this predicted power-law asymptotic behavior of *A*(*t*) with the exponent of −0.27, very close to the theoretical value of −0.25 (Fig. 3b). This slower power-law decay in these equal *A* + *B* systems compared to the *A* + *A* system arises because of spatial correlations that emerge after all closely adjacent *A, B* pairs have annihilated. This leaves only local regions of mostly *A* or mostly *B* particles, increasing the separation between reactant pairs. Because the analytical theory is derived in the continuum limit of an infinite domain, it is not exact for systems in a bounded domain or with excluded volume. Our simulations eventually diverge from the predicted power-law decay as the spatial correlations are restricted by our bounded domain length *L* (Fig 3b). As the mean displacement in 1D approaches the domain length, or 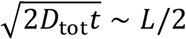, our discrete particles begin to feel the reflective boundary, accelerating the association compared to the continuum limit. We then see a return to faster decay of bounded reactants. To support this, we clearly see that our systems simulated with the same density but with a longer domain length *L* obey power-law relaxation dynamics for a longer time interval (Fig 3b).

### E. For reversible dimerization, volume exclusion of many-body systems will shift the expected equilibrium

The FPR algorithm is designed to ensure detailed balance for pairwise interactions. For reversible dimerization in a closed system, the many-body system *A* + *A* ⇌ *A*_2_ should relax to an equilibrium steady state where 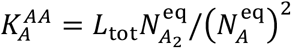 and mass conservation gives 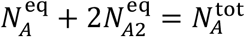. We find that even for density fractions of 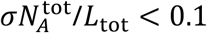, the space occupied by the *A* particles reduces the domain length sufficiently to concentrate the particles and reduce the fraction of unbound monomers in the simulations (Fig 4). Therefore, if we instead compare the predicted equilibrium using the accessible domain length *L* adjusted for volume exclusion using Eq. 10b, the simulations are in close agreement. Further, as the system becomes more dilute, the simulations and the predicted equilibria all converge to the same values, as expected (Fig 4). We verified that this result holds across parameter regimes spanning to diffusion-limited kinetics (Fig 4a-c), reinforcing that this equilibrium result is purely due to the effective reduction in domain size and not any numerical artifacts.

**Figure 4.**
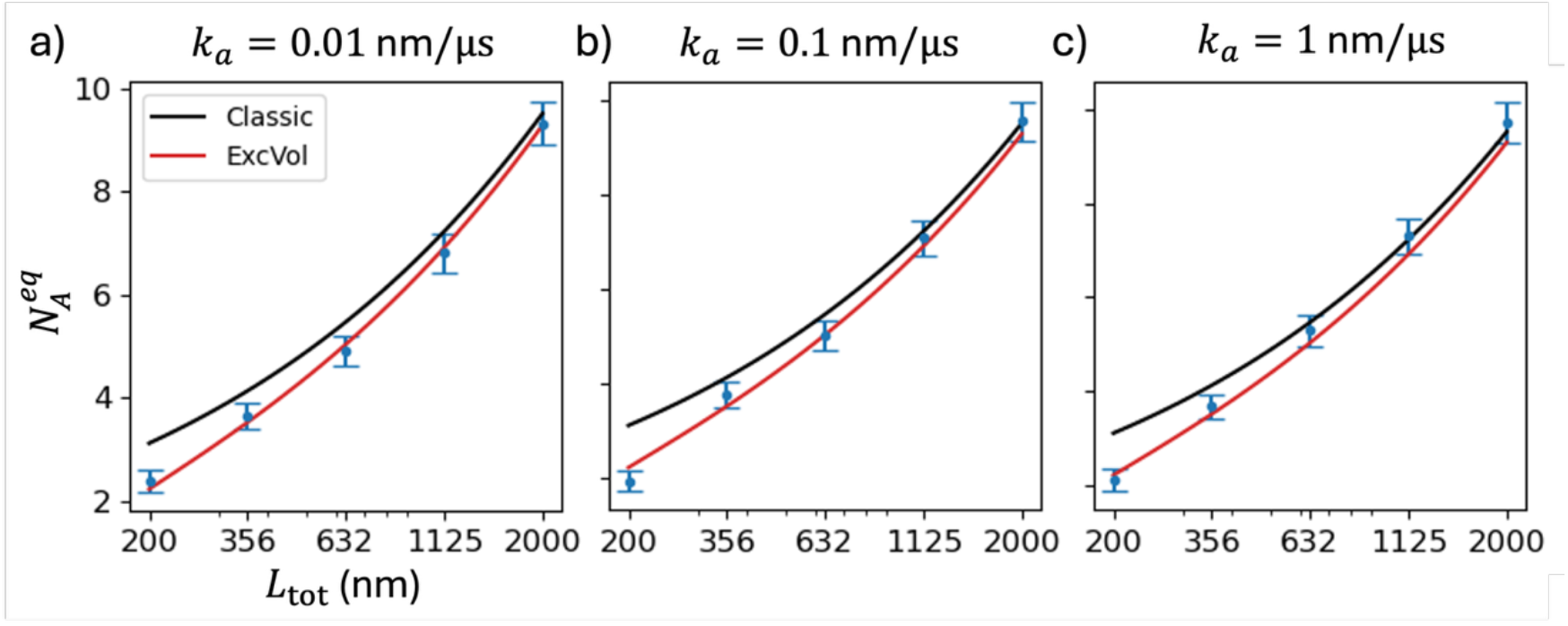
Equilibrium populations for many-body reversible dimer systems *A* + *A* ⇌ *A*_2_ are shifted by excluded volume. NERDSS simulations in blue points with SEM error bars. The black curve follows the theoretical equilibrium fraction of unbound monomers using the domain size *L*_tot_ (x-axis) whereas the red curve uses the accessible domain length *L* of Eq. 10, thereby predicting fewer free monomers. The association rates are a) *k*_*a*_ = 0.01 nm/*μ*s with *k*_*b*_ = 10 s^−1^ b) *k*_*a*_ = 0.1 nm/*μ*s with *k*_*b*_ = 100 s^−1^ as diffusion-influenced, and c) *k*_*a*_ = 1 nm/*μ*s with *k*_*b*_ = 1000 s^−1^ as diffusion-limited. Simulations use parameters: *Δt* = 0.025 *μ*s, *N*_*rep*_ = 48 repeats, *K*_*A*_ = 1000 nm, *D* = 0.2 *nm*^2^/*μ*s, *σ* = 1 *nm*, and 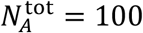.

### F. Kinetics of reversible dimerization for diffusion-influenced systems follows time-evolving rates

For the complete time relaxation from the initial monomeric state to the equilibrium state, we focused on systems at a moderate density where 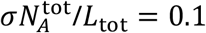 (Fig 5). We adjust both the ORE and MRE to use the accessible domain length *L* (Eq. 10). For the homodimerizing *A* + *A* ⇌ *A*_2_ simulations, we used *k*_*a*_ = 0.01 nm/*μ*s and a 10-fold slower diffusion coefficient of *D* = 0.01 nm^2^/*μ*s. As we revisit further in the next section, this places the kinetics in the strongly diffusion-influenced regime in 1D. In section II.C, we propose that to choose a single macroscopic rate constant for use in ORE, we use our derived density-dependent rate evaluated at the initial (most dense) monomer concentration, 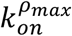 (Eq. 12). For these simulation conditions, we have 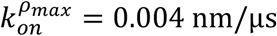, more than 2-fold slower than the intrinsic rate *k*_*a*_, because the purely diffusive rate to contact 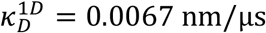 is slower than *k*_*a*_. We find that for three different dissociation rates with increasingly slower dissociation, the predicted kinetics using ORE with constant 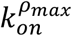 is initially too slow, and at longer times approaching equilibrium it is too fast (Fig 5a). The short-time deviation we observed as well in Fig 3a. The long-time deviation emerges as this single rate constant underestimates the growing separation between reactant pairs, predicting the significantly faster and exponential relaxation to equilibrium. However, using this rate 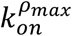 still presents what we consider an improvement over simply using *k*_*on*_ = *k*_*a*_, which describes only the shortest time dynamics well. With 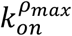 we more closely minimize the error between the ORE results and the RD dynamics and more accurately predict the correct half-time, *τ*_1/2_, to reach equilibrium, *A*(*τ*_1/2_) = *A*_0_/2.

**Figure 5.**
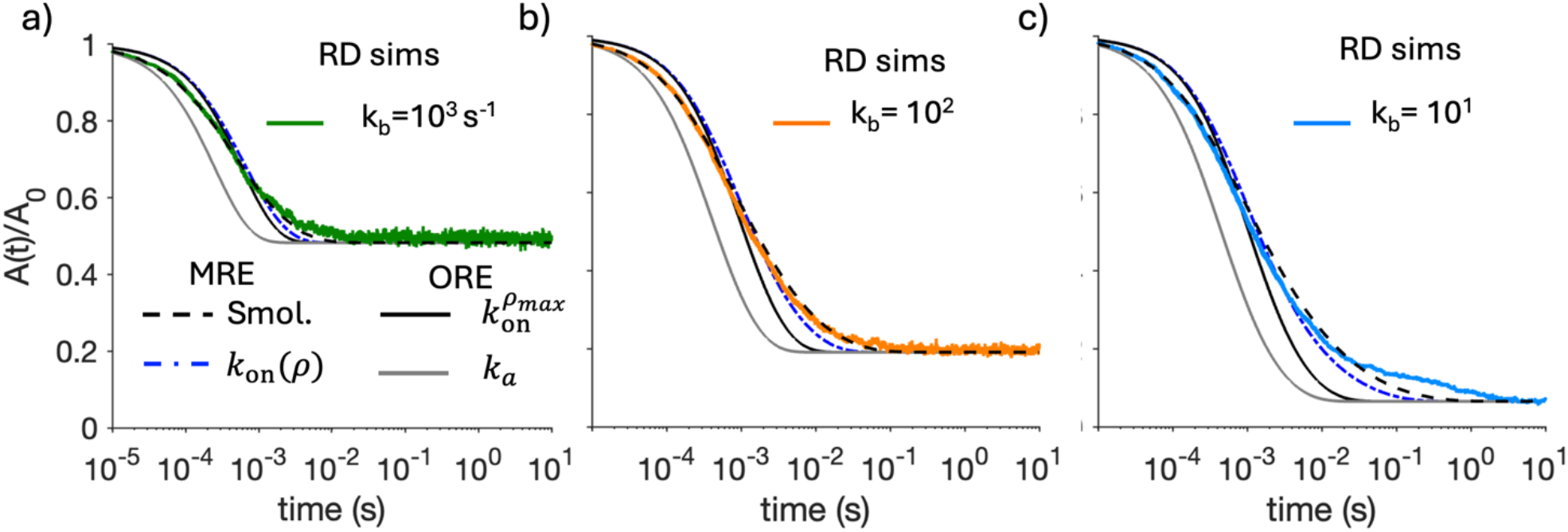
Reversible binding dynamics of strongly diffusion-influenced reactions follow time-evolving rates. The colored error-bar curves are NERDSS simulations with vertical *SEM*(*t*) bars for *A* + *A* ⇌ *A*_2_. A) *k*_*b*_ = 1000*s*^−1^ b) *k*_*b*_ = 100*s*^−1^c) *k*_*b*_ = 10*s*^−1^. We compare with 4 theoretical approximations. For the two MRE solutions, the Smoluchowski model 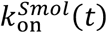 in black dashed agrees best in all cases, and the density-dependent rate 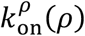 in blue dot-dash, shows the next-best agreement. The two ORE solutions use a constant 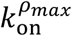 in solid black, whereas using *k*_*a*_ in gray shows the worst agreement. The rate equations all solve Eq. (23). The simulation parameters are 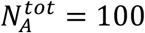, *σ* = 1 nm, *L*_tot_ = 1000 nm, *k*_*a*_ = 0.01 nm/*μ*s, *D*_*A*_ = 0.01 nm^2^/*μ*s. This results in *κ* = 1.5.

In contrast, we see that the MRE model with the time-dependent Smoluchowski rate produces very good agreement with the simulations, despite being an approximate theory, with deviations only emerging for long times with the slower dissociation rates (Fig 5c). Compared to the purely diffusion-limited irreversible association in Fig 3, here the barrier-limited rate *k*_*a*_ is orders-of-magnitude lower, although we still see that the simulations are slightly faster at intermediate times compared to the MRE, as observed in Fig 3. When we solve the MRE with our density-dependent rate 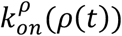, we find significantly better agreement than with a single rate constant, but the predicted short-time behavior is still too slow, similar to the results in Fig 3. At longest times the relaxation to equilibrium is a bit too fast.

### G. Excluded volume in systems with stable dimers and slow dissociation induce slower relaxation to equilibrium

The relaxation function to equilibrium is given by:

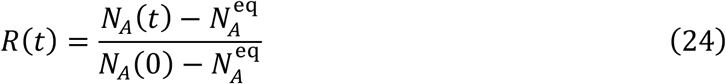

For fluctuating reaction-diffusion systems in 1D, this function relaxes with *t*^−1/2^ with pre-factors that depend on the concentrations, diffusion constants, and *K*_*A*_ of the system, assuming dilute point particles^27^. This contrasts with ORE, which have a well-known exponential relaxation to equilibrium, and contrasts with the MRE with the Smoluchowski model, which also fails to capture this power-law relaxation (despite predicting the *t*^−1/2^relaxation for *irreversible* systems), as shown in Fig. 6. The mean-field approximations inherent to the MRE rate models cannot fully capture the spatial correlations that emerge with the addition of dissociation events.

**Figure 6.**
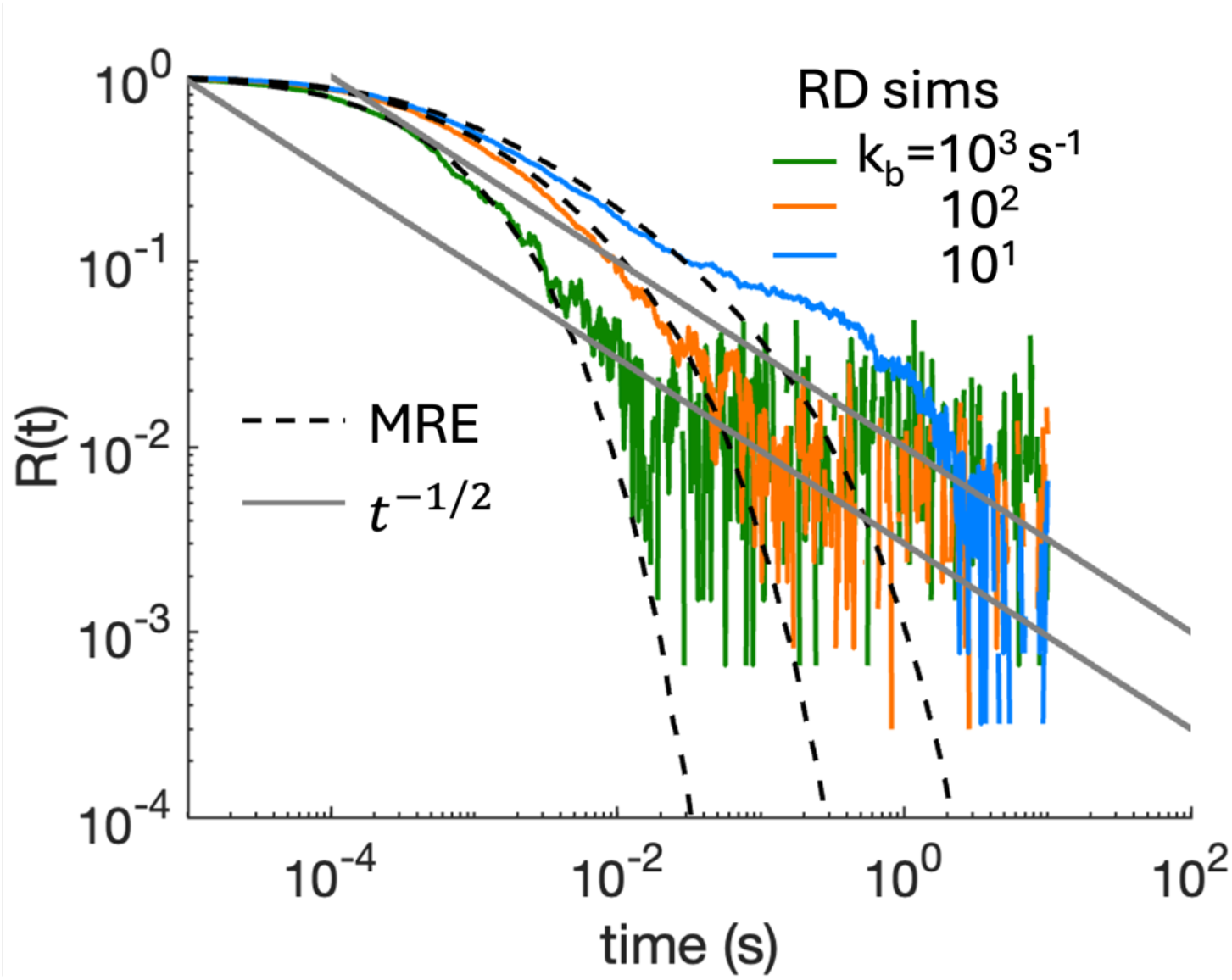
Relaxation to equilibrium in reversible systems can slow due to volume-excluding dimers. Using the same NERDSS data from Figure 5, we plot the relaxation to equilibrium Eq. 24. For comparison we show the MRE with the Smoluchowski rate model in black dashed, and the expected *ωt*^−1/2^ relaxation for point particles close to equilibrium. For green and orange, we see evidence in the RD simulations of the slower power-law relaxation to equilibrium, similar to that seen in 2D^37^. Prefactor *ω* = 3 and 10. For the most stable dimers in blue, we first see a visible plateau in the relaxation, as monomers are ‘blocked’ from association close to equilibrium by volume-excluding dimers.

Our simulation results in Fig 5 and Fig 6 demonstrate close agreement between the NERDSS simulations and the MRE with Smoluchowski rates at short to intermediate times, but the slower RD relaxation as the simulations approach equilibrium, as predicted^27^ (Fig 5c, Fig 6). Notably, we also observe for the slowest dissociation in Fig 5c, the relaxation kinetics of the RD simulations approaching equilibrium diverges further, first plateauing before relaxing to equilibrium. When the system forms stable dimers (high *K*_*A*_), we observe these long-time slow downs in the simulations compared to the *t*^−1/2^ limit of dilute systems (Fig 6). This emerges due to the physical barriers to association from volume-excluding *dimers* and is thus not captured in any of these theoretical models. We can anticipate these dimer blockade effects will be most pronounced in systems with strong spatial effects (or large *κ*), but also with a large *K*_*A*_, so equilibration is slowed. As spatial effects dominate the relaxation to equilibrium with slower equilibration, we find that so will the impact of the dimer blockades. Intuitively this makes sense as well for slow dissociation. If monomers are separated from a population of reactant partners by a dimer, dissociation is the only mechanism to allow them to find a partner. This effect is much less relevant for weak dimers, as a large pool of monomers is available, minimizing the disruption by dimer blockades. While this excluded volume effect is clearly visible in Fig 5c and Fig 6 (blue curve) the impact is still relatively modest.

### H. Reversible dynamics for distinct reactants *A* + *B* ⇌ AB will be sensitive to their linear arrangement

For the kinetics of distinct reactants, *A* + *B* ⇌ AB, we start by contrasting the relatively modest impact of excluded volume on the homodimer results above (Fig 6) with systems of many *A* and many *B* molecules that exclude volume with their bound states. Here we can expect dramatic changes in both the kinetics and the steady-state populations as they are fundamentally altered by an inability of reactants to ever bypass one another. Pairs of *A* and *B* reactants can be eternally partitioned away from one another depending on the initial conditions. In the extreme of partitioning all *B* molecules on the left and *A* on the right, only a single dimer could possible form due to excluded volume. While our NERDSS simulations can certainly capture this effect, one can already anticipate this result in principle. The emergence of these molecule blockades in real biological systems will be ameliorated by their ability to dissociate off the 1D filament and return to 3D.

To nonetheless assess the dynamics of reversible distinct reactions, we focus on the target problem with a single *A* particle and a diffusing population of *B* molecules randomly distributed on the filament, as only a single dimer can ever form (Fig 7). We again find that the ORE with the single rate 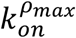 predicts overly slow association at short times, and overly fast association at long times compared to simulations (Fig 7a). We use the same density as above for the *B* reactants, 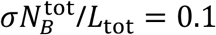 and adjust the domain length to the accessible length for all rate equations; previous results have shown that excluded volume between the diffusing *B* molecules does accelerate association kinetics^39,42^, and with *L* (Eq. 10) we account for that in our theoretical comparison. We found an important feature of this model is that the reactive flux into *A* is doubled when it is centered on the filament vs on the end (Fig 7a). At the center, the *B* molecules can approach from 2 sides, and thus the rate equations must incorporate a factor of 2 multiplying *k*_on_ for the centered *A* (Methods). As a result, these two models will reach a distinct equilibrium for the same value of *k*_*a*_ and *B* density. By doubling *k*_*a*_ for the end-of-filament target, we now recover the same equilibrium, albeit with noticeably slower kinetics (Fig. 7a). This is because doubling *k*_*a*_ is not equivalent to doubling the *k*_on_ that also accounts for diffusion (Eq. 11). For Fig 7a, because it has an excess of *B* reactants and only a single, stationary dimer state, we largely eliminate all many-body and excluded volume effects and therefore find excellent agreement between the MRE with the Smoluchowski rate and the NERDSS simulations. We also verified that allowing the target *A* to diffuse does not alter the equilibrium, but it does accelerate the kinetics for the end-of-filament target (Fig 7b). By diffusing from the end, the partners can be crowded into a smaller domain length, accelerating the association kinetics.

**Figure 7.**
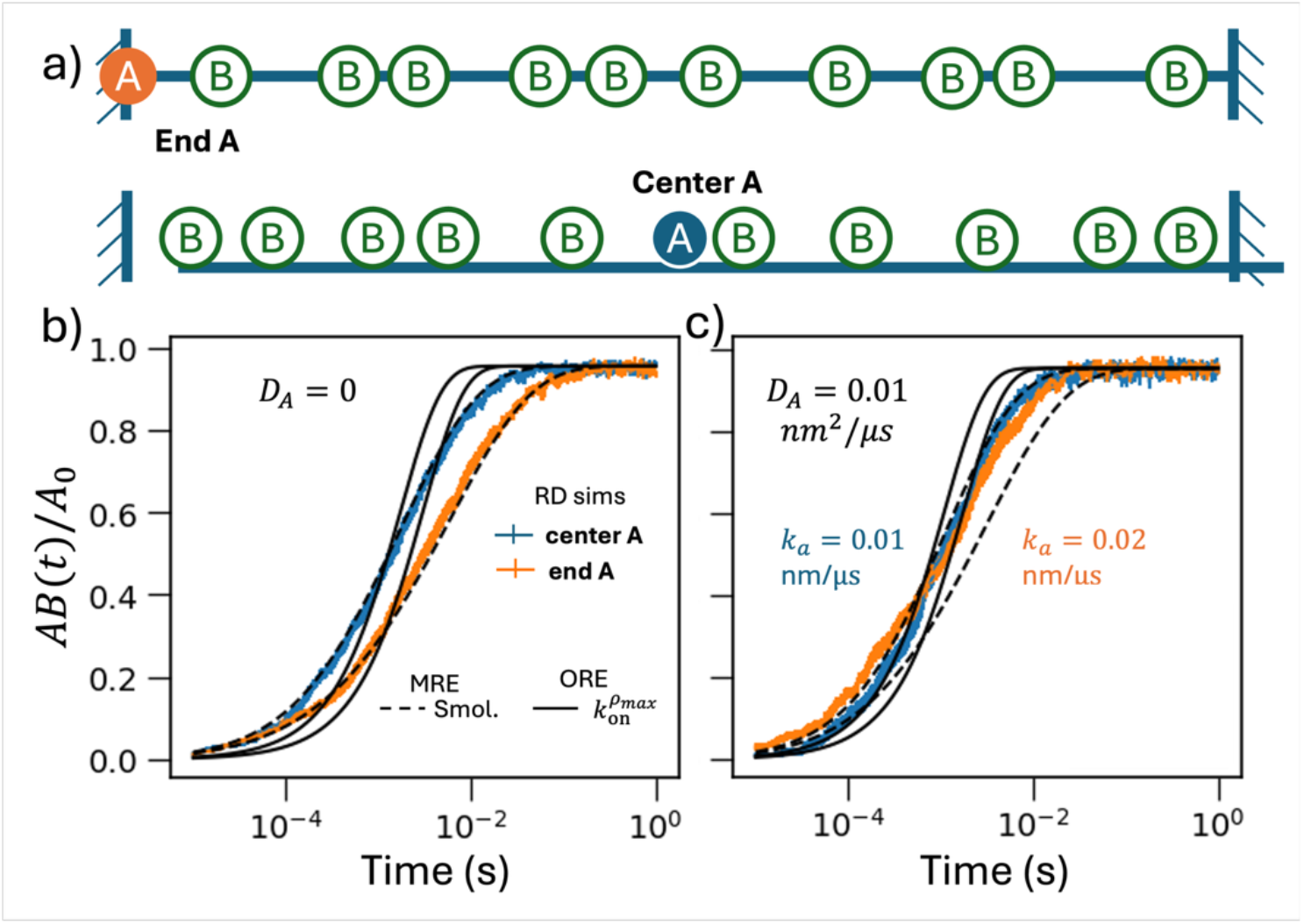
Reversible reactions with distinct reactants *A* + *B* ⇌ AB for a single *A* target particle. For simulations of reversible *A* + *B*, we use 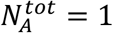 and 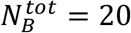 in *L*_tot_ = 200 nm. a) The target can either be placed at the end (orange A), or at the center (blue A), with the same density of B molecules. The flux through *A* is doubled when it is centered. B) The target *A* is stationary, *D*_*A*_ = 0 and *D*_*B*_ = 0.01 nm^2^/*μ*s. The reaction kinetics are noticeably slower when *A* is stationed at the end of the filament (orange) vs the center of the filament (blue curves) in the NERDSS simulations. For the center target, *k*_*a*_ = 0.01 nm/*μ*s, whereas for the end target, we increase *k*_*a*_ to 0.02 to achieve the same equilibrium state as the center target, but the kinetics are still slower. The MRE with Smoluchowski rate is in black-dashed, and the ORE is in solid black. c) The same system as (b), except now *A* can diffuse with *D*_*A*_ = 0.01 nm^2^/*μ*s. The center target has similar kinetics (blue curves), but the end target has noticeably faster kinetics now that *A* diffuses. Other simulation parameters are *σ* = 1 nm, and *k*_*b*_ = 100 s^−1^. Here *κ* = 1.5 for *k*_*a*_ = 0.01 nm/*μ*s and *κ* = 3 for *k*_*a*_ = 0.02 nm/*μ*s.

### I. Rate metric *κ* delineates regimes of rate-limited vs diffusion-limited dynamics in 1D

The results in Fig 5–Fig 7 used reaction parameters in the strongly diffusion-influenced regime (Table 1). These results illustrate that ORE with a single rate constant do not accurately capture the true RD kinetics of many-body association. However, we show here that we can predict rate-limited parameter regimes where the accuracy of a single rate constant becomes quite reliable within a reported error tolerance (Table 1 and Fig 9). In theory section II.4 we defined a ratio metric *κ* that quantifies the magnitude of the intrinsic rate *k* relative the diffusion-limited rate 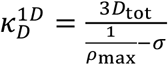. The diffusion-limited rate will change as the density changes, but overall we expect small *κ* to produce kinetics largely independent of *D* and *ρ*_max_, and thus well-defined by a single macroscopic rate, 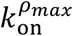. As we show in Fig. 8a, solving the ORE using 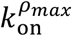 with *κ* =0.0032 produces very close agreement to the RD dynamics, as represented by the MRE solution which we use as a reliable proxy for the simulations. As *κ* increases to 0.032 and 0.32 (due to a larger *k*_*a*_) the simple ORE solution using 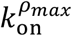 starts to deviate as quantified by the error Δ*A*(*t*), Eq. 21. The same trend is maintained for the reversible *A* + *B* target problem, although the error is shifted to the later stages of the reaction kinetics and is slightly higher for the target at the end (Fig 8b).

**Figure 8.**
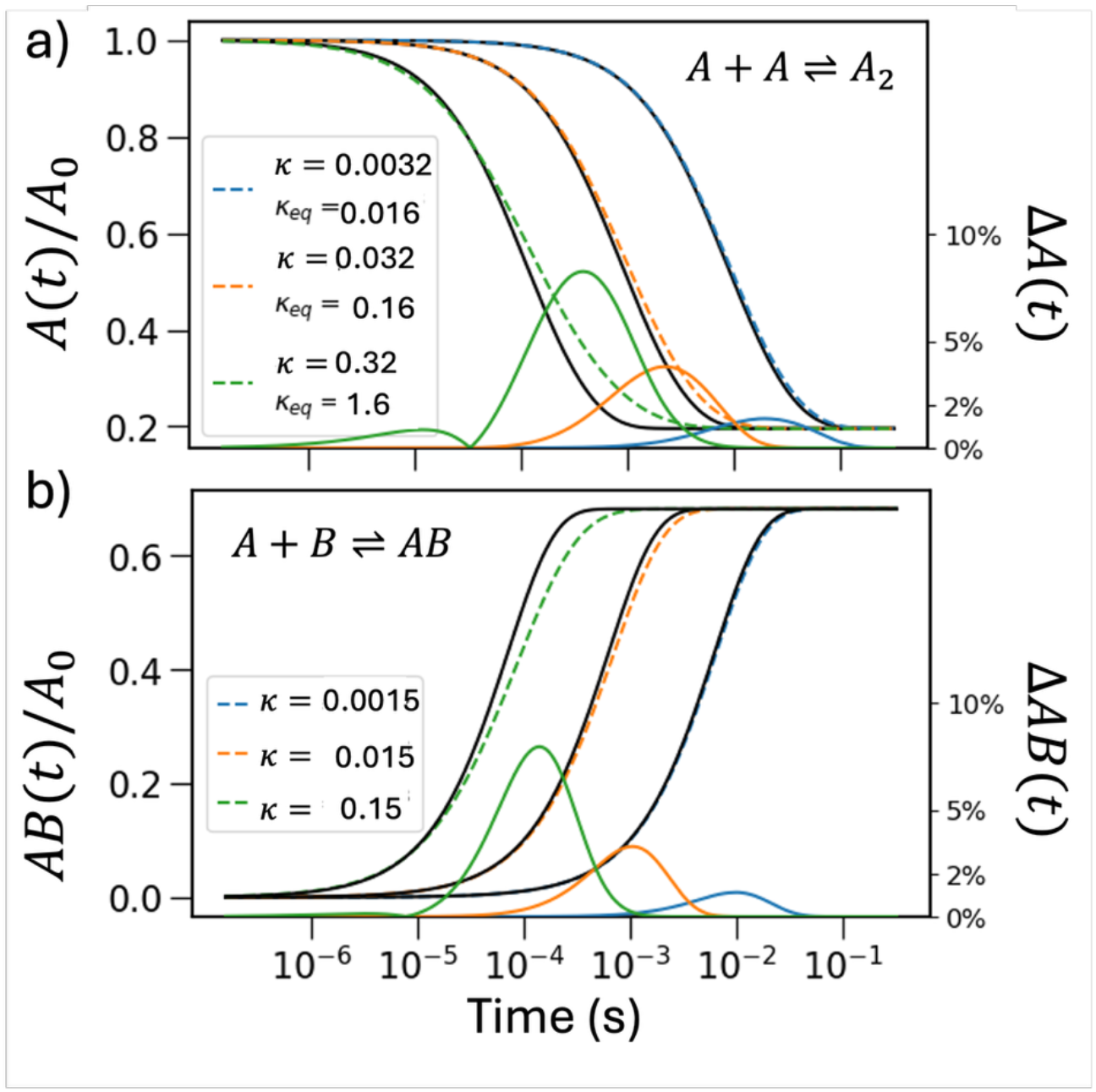
Comparison between single rate constant reaction kinetics and solutions to the MRE. Color dashed curves are the MRE solutions with the rate model from 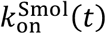 as a good approximator of the RD dynamics, and black curves are ORE solutions with 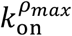. The color solid lines are normalized absolute error between two curves (see Methods, Eq. 21). a) Homodimerization with 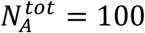, *σ* = 1 nm, *L*_tot_ = 2000 nm, *D*_tot_ = 2 nm^2^/*μ*s, and *K*_*A*_ = 200 nm. *k*_*a*_ = 0.001, 0.01, and 0.1 nm/*μ*s for the blue, orange, and green curves, producing *κ* values of 0.0032, 0.032, and 0.32. *k*_*b*_ = *k*_*a*_/*K*_*A*_ accordingly. b) Target searching with target-at-end scenario, with 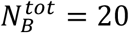, *σ* = 1 nm, *L*_tot_ = 200 nm, *D*_*B*_ = 2 nm^2^/*μ*s, and *K*_*A*_ = 20 nm. *k*_*a*_ = 0.001, 0.01, and 0.1 nm/*μ*s for the blue, orange, and green curves, producing *κ* = 0.0015, 0.015, 0.15. *k*_*b*_ = *k*_*a*_/*K*_*A*_.

To establish useful regimes of *κ*, we combine theoretical analysis of the Smoluchowski model with empirical measures of the error, max (Δ*A*(*t*)). When the rate from the Smoluchowski model 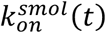 is still within a factor of (1 − *δ*)*k* where *δ* is small, we expect good agreement with a single macroscopic rate. We therefore define this function used for plots in Fig. 9:

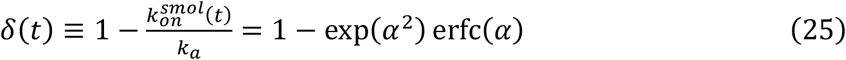

**Figure 9.**
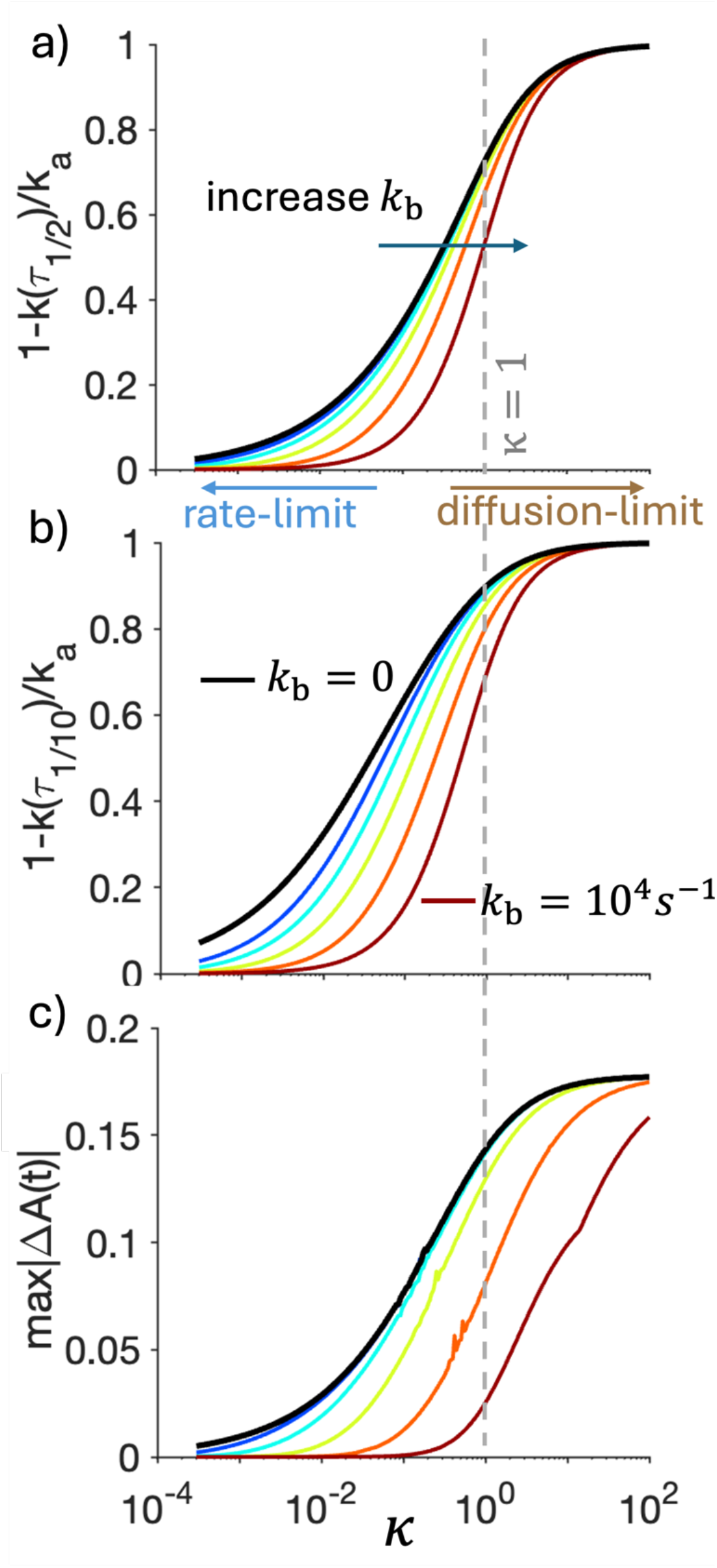
Increasing values of *κ* produce monotonically increasing deviation from kinetics predicted by a single rate constant. All three subplots have *κ* along the x-axis, gray dashed line is *κ* = 1. Each colored curve predicts *κ* dependence with an increasing dissociation rate from left to right, from *k*_*b*_ = 0 (Black curve) to *k*_*b*_ =1 (blue), 10 (cyan), 100 (green) and 1000 (orange) and 10^4^s^−1^ (maroon curve). The deviations decrease as *k*_*b*_ increases. A) our function *δ*(*t*) defined in Eq. 25 predicts how strongly the rate will have slowed at *t* = *τ*_1/2_, with small values indicating a single rate constant is accurate, and large values indicating strong density-dependence of the rate. At *κ* = 1, the rate will slow by ~75% for irreversible systems at *τ*_1/2_. B) same as (a) but evaluated at 90% completion, shifting to larger *δ*. C) The absolute error max|Δ*A*(*t*)| Eq. 21. The error plateaus at large *κ*, as solutions converge to diffusion-limited. Simulations were run with *A*_0_ = 0.1/*nm, σ* = 1*nm, D*_tot_ = 0.1*nm*^2^/*μs*, and *κ* is increased via *k*_*a*_. Results are nearly density and *D*_tot_ independent for (a-b).

Where from Eq. 6 the argument 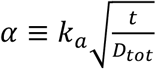 and *δ* ∈ [0,1]. We have for small rate changes, or *δ* ≤ 0.1, that *α* ≤ 0.1. For a maximally 2-fold slow-down in the rate, or *δ* ≤ 0.5, then *α* ≤ 0.77, whereas we will see a 10-fold or larger slow-down in the rate when *α* ≥ 5.5 To use this result where *α* controls rate behavior requires the definition of a characteristic timescale *τ*, which all our results above demonstrate should clearly be dependent on the 1D particle density *ρ*. We use the timescale to proceed to a fraction *s* of the initial relative to steady-state concentration via *τ*_*s*_, as the solution to *A*(*t* = *τ*_*s*_) − *A*_*eq*_ = *s*(*A*(0) − *A*_*eq*_), where 0 < *s* ≤ 1. We plot *δ*(*τ*_1/2_) or 50% completion in Fig 9a and 90% completion *δ*(*τ*_1/10_) in Fig 9b. To define simpler analytical expressions for *τ* (and therefore *δ*(*τ*)) that retain dependence on density and diffusion, we use the analytical solution to the ORE with 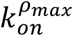. For irreversible systems of *A* + *A, A*_*eq*_ = 0, we have 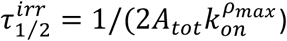, and 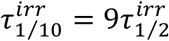. At 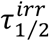, we have that 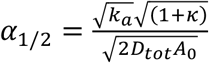, as shown by the black curve in Fig. 9a. For reversible systems we have: 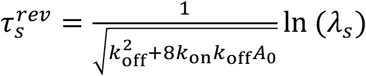, where 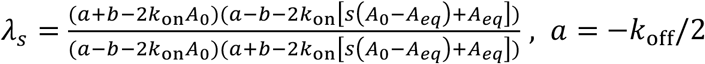, and 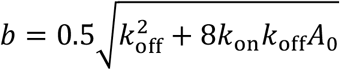. We note that the expected deviation between *k*_*a*_ and 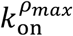 can be compactly predicted via: 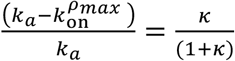.

In Fig. 9, we demonstrate how with increasing *κ*, we predict monotonic deviations from the assumption of a single rate constant. In Fig 9a-b we illustrate how as the reaction proceeds to halfway completion, the rate of association will have slowed from *k*_*a*_ by only ~10% for *κ* = 0.01, but by ~75% for *κ* = 1. Naturally, the rate has changed more by the time we reach *τ*_1/10_ in Fig 9b. For the same *κ* value, the largest error occurs for *k*_*b*_ → 0, as these irreversible processes take the longest to reach steady-state. With more rapid dissociation, the system also does not reach the lowest monomer densities. These results are effectively independent of variations in the density or particle diffusivity. Finally in Fig 9c, we show the error incurred between the kinetics predicted by ORE with a single rate constant 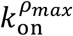, vs the MRE solution with *k*_*smol*_(*t*) as the RD proxy. Because 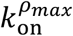 generates relaxation that is too slow for the first half and too fast for the second half, it tends to minimize the overall error, generating absolute error limited to <20% of *A*_0_. We note that with lower *A*_0_, the error is slightly smaller, whereas at *A*_0_ = 0.5, the max error plateaus at ~23%, so it is density dependent. These quantitative results are overall conserved for the reversible *A* + *B* target problem with centered *A*. With the *A* at the end, however, the max error metric (Fig. 9c) is ~35% worse, due to the slower relaxation for this spatially asymmetric set-up.

Based on these theoretical results and the empirical error in relaxation kinetics, we propose regimes of *κ* for rate-limited through to diffusion-limited reaction dynamics (Table 1). We will use theory and simulations at the irreversible limit, as the ‘worst-case’ scenario (Fig 9), noting that clearly one gets better agreement for systems with frequent dissociation (Fig 9c). For *κ* < 0.01, we see the rate at halftime completion will still be within 13% of *k*_*a*_, and the full time-resolved kinetics will have <3% error, which emerges as steady-state is approached. Even at the worst-case, we propose this as rate-limited. For more stringent error tolerance, an even lower *κ* is required. For 0.01 < *κ* < 0.1, we see the influence of diffusion, producing 35% drop in the rate by halftime, and 65% by *τ*_1/10_, with 8% error in many-body relaxation, so we propose this as diffusion influenced. From 0.1 < *κ* < 1, we see strong diffusion influence, with half-time rates that are 2-fold slowed, and >10% error. As *κ* > 1, and particularly for *κ* > 10, we see convergence to the purely diffusion limit. While we recommend 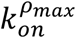 as a mean-field macroscopic rate for continuum rate models, for *κ* > 0.1 and certainly *κ* > 1, the kinetics shows clear density dependence of the association rate as the reaction proceeds. These conclusions are compactly presented in Table 1.

## V. DISCUSSION and CONCLUSIONS

In 1D, the kinetics of molecular association and dissociation can be highly sensitive to the details of the model for parameters in the diffusion-influenced regime. Excluded volume of monomers and dimers, the reversibility of components, and for distinct components, their linear arrangement, will all impact the kinetics and even the steady-state. However, despite this sensitivity, we here identified regimes of system density and reaction parameters where the 1D kinetics is well-approximated by ordinary rate equations (ORE) with a single rate constant. The success of this coarse-grained single-rate approach will depend on the reaction parameters, which specify the value of *κ*. We provide guidelines for model selection and expected error in Table 1. Single rate constants are not only used in ORE, but also continuum RD models (which lack noise or discrete particle resolution), and the RDME framework^25^. In all our simulations above, for example, the initial conditions were homogeneous or ‘well-mixed’, which for continuum RD models generates the same relaxation as the ORE models, demanding a macroscopic rate *k*_on_ and not *k*_a_. While RDME captures discrete particle numbers, it does not resolve any spatial correlations that emerge below the lattice spacing, and hence similarly requires a macroscopic rate dependent on this lattice spacing^51^. Here we demonstrate that this rate be defined relative to the density of reactants, where to improve agreement with the stochastic particle-based dynamics, the rate should update as the local density changes. Our density-dependent rates (Eq. 11) can thus locally respond to both decreases and increases in reactant density when the 1D reactions are coupled to other biochemical reactions or localization from 3D, which is not possible when using the Smoluchowski rate model directly. For the active field of ‘whole-cell’ models^52–55^, binding rates are critical parameters to achieve cell-scale spatio-temporal dynamics, and with increasingly detailed models of assembly and localization to DNA, inclusion of 1D reaction kinetics will be necessary to reproduce facilitated diffusion^56^. While using a single rate constant in 1D is an approximation that will generate deviations in the continuum kinetics compared to the ‘true’ particle-based kinetics, we summarize in Table 1 and Fig 9 that it can produce close agreement across models for *κ* ≪ 1.

For the expected regimes of *κ*, we expect the full range of values to be accessible by proteins sliding on DNA. The diffusion coefficients *D* of nonspecifically bound proteins sliding along DNA are measured in ranges of ~0.001-1 um^2^/s^56^. The relevant lengthscales of DNA segments in eukaryotes is 60nm up to ~1000 nm due to nucleosome barriers^17,57^. Although in prokaryotes, proteins can in principal slide the length of the genome, the 1D sliding trajectories are expected to be roughly 50-200 nm^9^. For a pair of proteins, we therefore expect densities of 0.002-0.05 /nm, consistent with the systems studied here. The values of *k*_a_ can vary the widest, with the diffusion limit (*k*_a_ → ∞) likely more common for protein-DNA binding, where for a pair of particles the effective macroscopic rate then scales linearly with density and *D*. For protein-protein interactions^58^ and even protein-DNA target binding, collisions between partners are not always reactive^59^, and there is no simple formula to predict 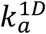 from 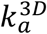, beyond stating that 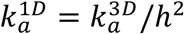, where *h* is a lengthscale that encodes changes to the binding energy landscape and barrier crossing times. Practically, however, these rates can be extracted from molecular dynamics simulations, as has been done for 2D rates^60^ and 2D equilibrium constants^61,62^, and they could be inferred from experimental single-molecule data of proteins sliding and binding while DNA-bound^17^. Given the diversity of transcription factors and their combinatorial contacts with one another^63^, we speculate that *k*_a_ can span to the slower association values required for small *κ*, supporting 1D reaction kinetics from the rate-limited through to diffusion limited.

A primary limitation of characterizing kinetics of association and dissociation between monomers using this particle-based reaction-diffusion framework is that we have not explicitly included all physical features of protein dynamics on DNA. DNA has two strands, and bypassing is therefore possible on this two-way ‘road’ if the contact between the protein and the DNA does not span both strands. Stability on nonspecific sequences is not fully sequence-independent: repeat sequences can be more stabilizing than fully random sequences^64^, which could create heterogeneous diffusion along DNA. DNA mechanics, whether due to supercoiling or topological domains or nucleosome positioning, can locally alter binding affinities^65^. Ultimately, however, building models that can reach the time and length-scales of single-molecule tracking experiments^66^ demands coarse-graining out the full atomistic details. Even with the addition of DNA structure and dynamics^67^, it will nonetheless be important to allow proteins to diffuse and react along the polymer backbone. The modeling paradigm used here offers a resolution that is therefore important for bridging molecular to cell-scale dynamics.

Overall, the modern biophysical characterization of macromolecular interactions on effectively 1D filaments like DNA reinforces the important role that 1D diffusion plays in controlling binding specificity and selectivity for DNA targets throughout the genome^14,17,59^. While in vitro and theoretical studies of binding affinities and kinetics more commonly have focused on monomers^14,17,59,64^, proteins like the GAGA factor form dimers and higher-order assemblies that bind and bridge chromatin segments^68^, and their formation and stability will depend, in part, on their 1D diffusion and interactions^17,19^. The theoretical results and stochastic, particle-resolved simulations presented here, which capture volume exclusion as it impacts dynamics, integrate directly into more complex models and environments such as these, including the reversible localization of proteins from 3D to 1D DNA ^19^ and higher-order protein oligomerization^69,70^. This work will thus help facilitate quantitative models for interpretable and predictive dynamics of coarse-grained molecular dynamics in the cell.

## ACKNOWLEDGEMENTS

We gratefully acknowledge support from an NIH grant U01DK127432 to MEJ, and an NIH grant R35GM161901 to MEJ.

## AUTHOR DECLARATIONS

Conflict of Interest: The authors have no conflicts to disclose.

## DATA AVAILABILITY

The data that support the findings of this study are available within the article and a github repository: https://github.com/JohnsonBiophysicsLab/densityVolume1D. All software is available open source at https://github.com/JohnsonBiophysicsLab/NERDSS.

